# MFN2-dependent recruitment of ATAT1 coordinates mitochondria motility with α-tubulin acetylation and is disrupted in CMT2A

**DOI:** 10.1101/2023.03.15.532838

**Authors:** A. Kumar, D. Larrea, M.E. Pero, P. Infante, M. Conenna, G.J. Shin, W. B. Grueber, L. Di Marcotullio, E. Area-Gomez, F. Bartolini

## Abstract

Acetylated microtubules play key roles in the regulation of mitochondria dynamics. It has however remained unknown if the machinery controlling mitochondria dynamics functionally interacts with the α-tubulin acetylation cycle. Mitofusin-2 (MFN2), a large GTPase residing in the mitochondrial outer membrane and mutated in Charcot-Marie-Tooth type 2 disease (CMT2A), is a regulator of mitochondrial fusion, transport and tethering with the endoplasmic reticulum. The role of MFN2 in regulating mitochondrial transport has however remained elusive. Here we show that mitochondrial contacts with microtubules are sites of α-tubulin acetylation, which occurs through the MFN2-mediated recruitment of α-tubulin acetyltransferase 1 (ATAT1). We discover that this activity is critical for MFN2-dependent regulation of mitochondria transport, and that axonal degeneration caused by CMT2A MFN2 associated mutations, R94W and T105M, may depend on the inability to release ATAT1 at sites of mitochondrial contacts with microtubules. Our findings reveal a function for mitochondria in regulating acetylated α-tubulin and suggest that disruption of the tubulin acetylation cycle play a pathogenic role in the onset of MFN2-dependent CMT2A.

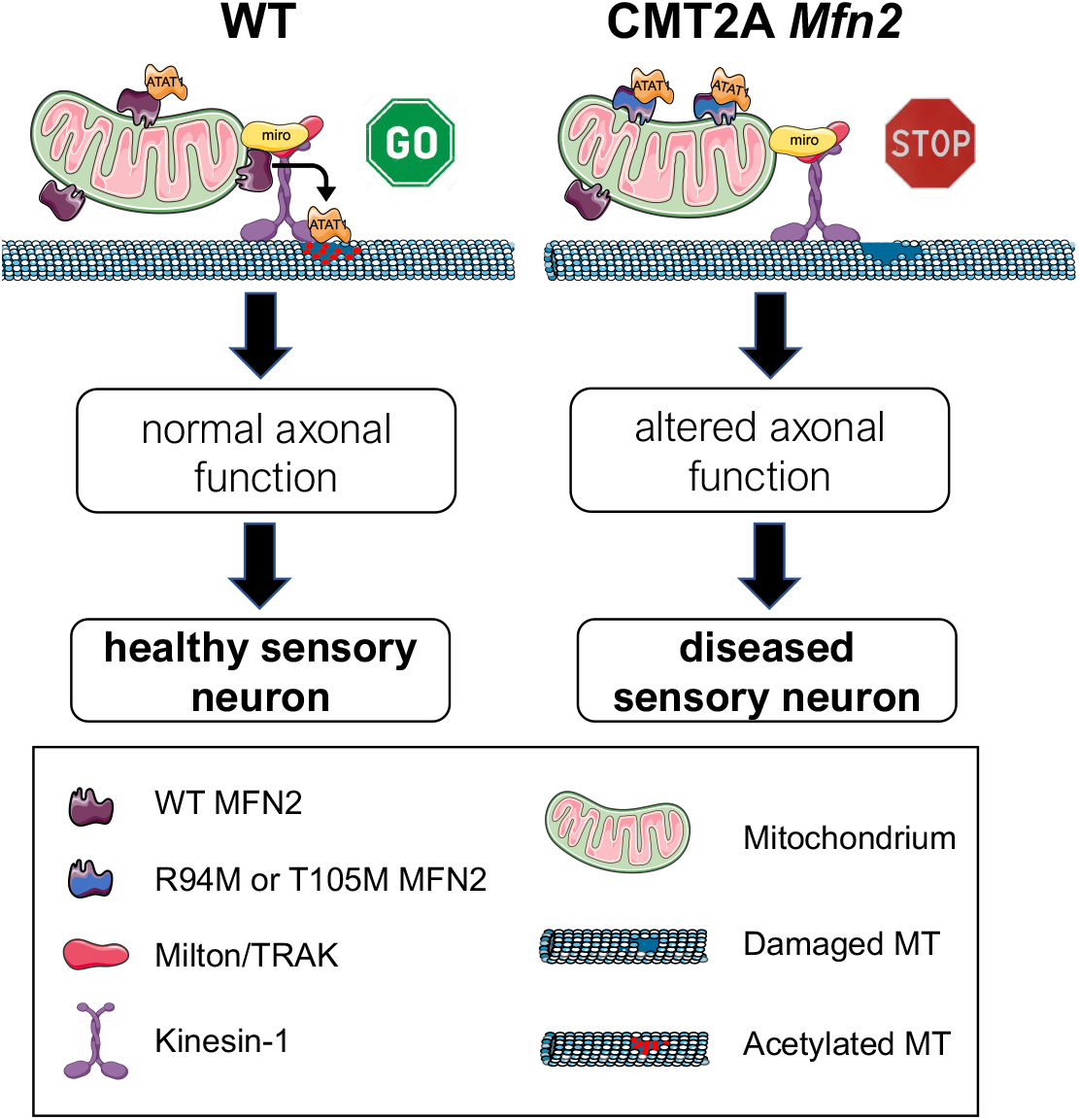

**Highlights:**

- Mitochondria contacts with MTs are hotspots of α-tubulin acetylation through the recruitment of ATAT1 by MFN2
- Mutations in MFN2 associated with CMT2A disease lose this activity by sequestering ATAT1
- Distal axonal degeneration caused by loss of MFN2 depends on acetylated tubulin-mediated mitochondria transport

**eTOC:** Recruitment of ATAT1 to mitochondria by MFN2 is critical for axonal viability through the regulation of mitochondria transport, and is disrupted in CMT2A

## Introduction

Microtubules (MTs) are key cytoskeletal elements involved in a multitude of functions in all eukaryotic cells. MTs are polarized polymers (comprised of a plus and minus end) constructed by the regulated polymerization of α- and β-tubulin dimers, and their dynamic nature allows them to switch between growth and shrinkage [1]. In neurons, MTs are critical because they support trafficking while providing segregation of functional sub compartments [2]. In vertebrates, MT plus ends uniformly orient toward the distal end of axons, while MTs are arranged with mixed polarity in dendrites [3]. In addition to ensuring structural support, MTs act as intracellular highways for protein motors of the kinesin and dynein family to deliver cargoes via anterograde or retrograde transport respectively. Directional transport is enabled by the structural polarity of MTs, which is recognized by motor proteins that drive transport to either the minus end (dynein) or plus end (most kinesins) [4]. In addition to structural polarity, tubulin post-translational modifications (PTMs) that accumulate on stable MTs are crucial modulators of neuronal transport and their dysregulation has been recently associated with the pathogenesis of both neurodevelopmental and neurodegenerative diseases [5, 6].

Acetylated MTs are a subset of stable MTs with key roles in the regulation of axonal transport and mitochondrial dynamics: along with providing preferential tracks for kinesin-1- and dynein-dependent mitochondria transport, two properties of acetylated MTs, their stability and flexibility, make them uniquely adapted to sustain mechanical stress caused by surface tension and organelle/organelle interactions [7–9]. It is perhaps thanks to these properties that mitochondria fusion, fission and ER/mitochondria contact occur selectively on acetylated MTs [10, 11].

Acetylation of lys-40 in α-tubulin is predominantly regulated by the tubulin N-acetyltransferase 1 (αTAT1 or ATAT1) and histone deacetylase 6 (HDAC6), two soluble enzymes that catalyze the forward and backward reaction, respectively [12–15]. Tubulin acetylation on lys-40 is an a-tubulin PTM marking the luminal surface of MTs [16, 17], a unique feature that helps the MT lattice cope with mechanical stress via reduced lateral interactions between protofilaments [7] and presumably by facilitating MT self-repair through incorporation of GTP-bound tubulin subunits [9, 18].

It is thought that in neurons a large pool of ATAT1 is recruited to the MT lattice by vesicle “hitchhiking” prior to entering the lumen at MT ends or at cracks in the MT lattice [19–22]. In migrating cells, clathrin-coated pits control MT acetylation through a direct interaction of the ATAT1 with the clathrin adaptor AP2 [23]. The rules dictating the selection of the docking sites on vesicles, clathrin-coated pits or MTs are however unknown. Similarly unexplored is whether other organelles can act as docking sites for ATAT1 binding. These are important questions to address, as either hypoacetylation or hyperacetylation of tubulin are predicted to negatively affect MT-dependent functions, either by reducing the local flexibility of the MT (thus promoting further breakage upon bending) or by inhibiting MT dynamics while promoting premature tubulin longevity.

In sensory neurons acetylated tubulin is an essential component of the mammalian mechanotransduction machinery through its regulation of cellular stiffness and TRP channel activity [24–26], and loss of acetylated tubulin was reported as a neuropathological feature of vincristine-induced toxicity [27]. Indeed, enhancing tubulin acetylation by HDAC6 inhibitors has been largely successful in restoring axonal integrity and myelination of toxic and genetically inherited forms of peripheral neuropathy, prompting multiple companies to develop and test HDAC6 inhibitors in models of peripheral neuropathies including Charcot-Marie-Tooth disease (CMT) [27–29]. Given the multitude of HDAC6 substrates in addition to tubulins, however, the mechanisms underlying this rescue remain unclear.

Charcot-Marie-Tooth type 2A (CMT2A) disease is a predominantly axonal form of familial peripheral neuropathy causing sensory loss that results from degeneration of long peripheral axons [30]. Inherited dominant mutations in the mitochondrial fusion protein mitofusin-2 (MFN2), a large GTPase residing in the outer mitochondrial membrane (OMM) and endoplasmic reticulum (ER), are the most common causes of CMT2A, and the majority of MFN2 mutations affect the amino terminal GTPase domain, with disease onset in the first two years of life and an aggressive clinical course [31, 32]. Like acetylated tubulin, MFN2 plays crucial roles in mitochondria dynamics, including regulation of fusion, motility, and ER/mitochondria contacts [33]. Defects in mitochondria dynamics are typically associated with CMT pathogenesis, including CMT caused by mutant MFN2 [34–37]. However, the mechanism by which mutant MFN2 contributes to CMT2A remains elusive.

Together with mitofusin-1 (MFN1), MFN2 regulates mitochondrial fusion, which is essential to maintain proper mitochondrial distribution, shape and degradation [33]. In addition, MFN2 plays a critical role in ER-mitochondrial tethering, which is independent of its fusion function, by bonding mitochondria with mitochondria-associated ER membranes (MAMs) to allow for ATP, Ca^2+^ and lipid transfer [34, 38–41]. Accordingly, MFN2 also regulates the rate of cholesterol esterification, which was proposed to be a proxy of MAM function, and the extent of contact points between the ER and mitochondria [42].

A fusion-independent role for MFN2 in regulating mitochondrial axonal transport has been reported. Loss of MFN2 or MFN2 disease mutants selectively alter mitochondrial axonal motility and distribution [35, 43, 44]. In addition, MFN2 deficiency in human spinal motor neurons interferes with mitochondrial transport while reducing both mRNA and protein levels of kinesin and dynein motors [45], which may further contribute to impaired mitochondrial motility. Furthermore, both MFN1 and MFN2 interact with mammalian miro (miro1/miro2) and milton/TRAK (OIP106/GRIF1) proteins, members of the molecular machinery that links mitochondria to kinesin motors [43], and overexpression of MFN1 rescues the axonal degeneration caused by MFN2 mutants *in vitro* and *in vivo* [44, 46].

Altogether, these data support the notion that MFN2 may directly influence mitochondrial positioning, and that loss of this function contributes to the degeneration of long axons, which are particularly sensitive to failures in meeting local energy demands. This model is consistent with the observation that most of the genes mutated in predominantly axonal forms of CMT have roles in mitochondrial motility, suggesting that impaired mitochondrial transport may be a common mechanism of pathogenesis shared by seemingly unrelated proteins associated with an increased risk for CMT [36]. Despite this compelling evidence, the molecular basis underlying MFN2-dependent regulation of mitochondria positioning remains poorly understood. Furthermore, while loss of MFN2 induces axonal neuropathy, the detailed mechanisms by which MFN2 deficiency results in axonal degeneration are unestablished.

In this study, we find that mitochondria contacts with MTs are hotspots of tubulin acetylation through the recruitment of ATAT1 by MFN2 onto mitochondrial outer membranes and that this activity is affected by MFN2 R94W and T105M CMT2A mutations. Furthermore, we provide evidence that axonal degeneration caused by MFN2 loss of function in DRG neurons depends on loss of acetylated tubulin by disrupting mitochondria motility rather than mitochondrial fusion or tethering with the ER.

## Results

### MFN2 is a novel regulator of tubulin acetylation

Despite the intimate relationship between acetylated MTs and mitochondria dynamics, it has remained unknown if the machinery controlling mitochondria motility and/or hetero-homotypic mitochondrial contacts functionally interacts with the α-tubulin acetylation cycle. To test this hypothesis, we measured levels of acetylated α-tubulin in immortalized Mfn2 KO mouse embryonic fibroblast (MEF) cells with reported defects in mitochondria dynamics and functional tethering with the ER [47]. By immunoblot and immunofluorescence analyses, we found that while de-tyrosinated tubulin levels were unaffected, loss of MFN2 reduced acetylated tubulin by more than 50%compared to WT controls (Fig. 1A-E and Fig. S1A-D). Loss of acetylated tubulin in these cells also correlated with a decrease in the abundance of GTP-tubulin islands, the putative entry sites for the tubulin acetyltransferase ATAT1 into the MT lumen and hotspots of MT self-repair by incorporation of GTP-bound tubulin subunits [48–51] (Fig. 1F,G).

**Figure 1.**
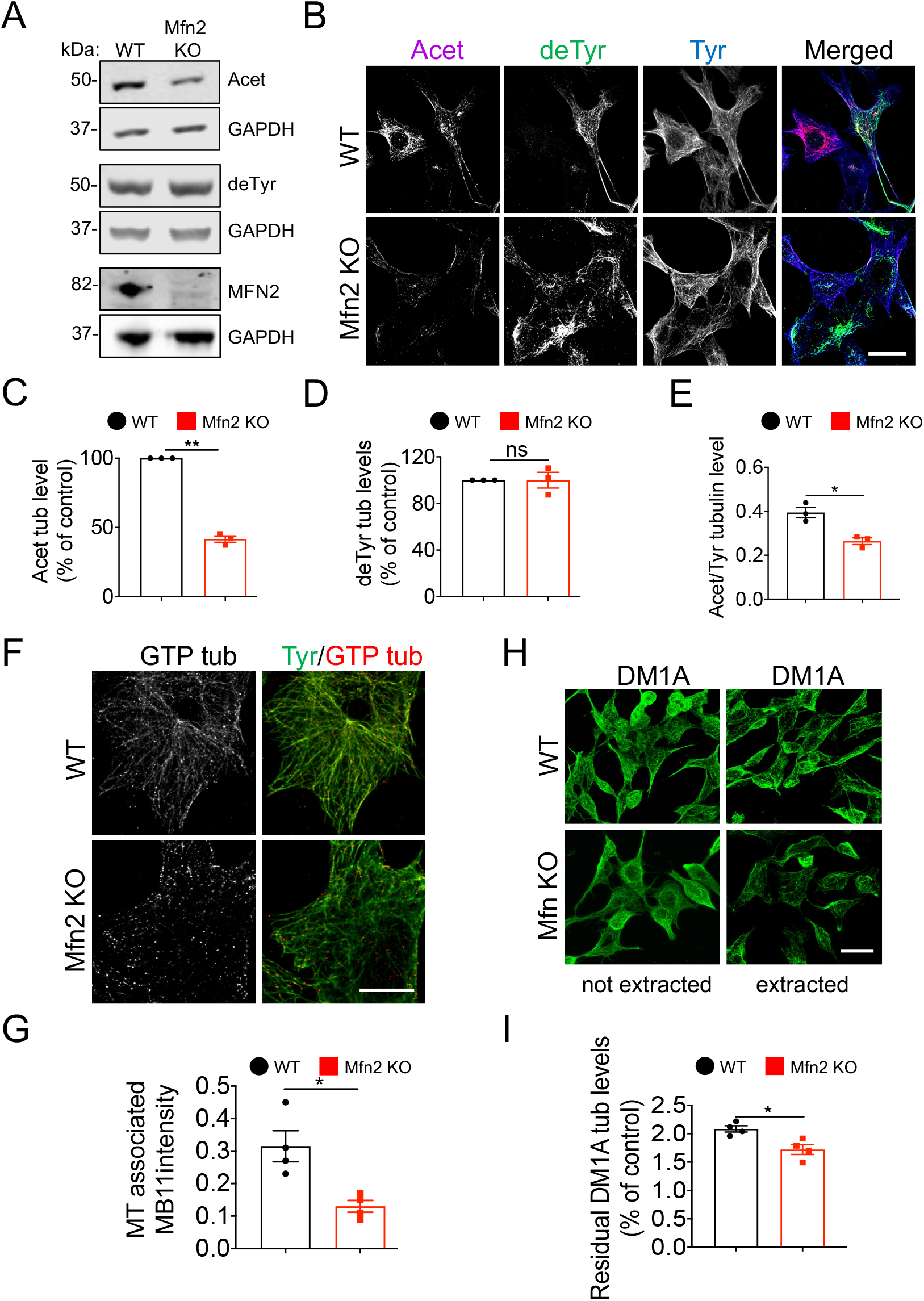
MNF2 regulates α-tubulin acetylation, MT dynamics and MT stability in MEFs. (A) Representative immunoblot of WT and Mfn2 KO whole MEF lysates. Acet, acetylated tubulin; deTyr, detyrosinated tubulin; Mfn2, mitofusin 2; GAPDH, loading control. (B) Representative immunofluorescence images (max projections from z-stacks) of WT and Mfn2 KO MEFs. Tyr, tyrosinated tubulin; Acet, acetylated tubulin; Tyr, tyrosinated tubulin. (C) Quantification of acetylated (Acet) and (D) detyrosinated (deTyr) tubulin signal normalized to WT control levels (n=150-175 cells). (E) Quantification of acetylated (Acet) to tyrosinated (Tyr) tubulin immunofluorescence signal intensity ratio in WT and Mfn2 KO MEFs. (F) GTP tubulin staining in WT and Mfn2 KO MEFs using hMB11 antibody staining. (G) Quantification of GTP-tubulin (MB11) immunofluorescence signal intensity associated with MTs in WT and Mfn2 KO MEFs (n=20-25 cells). (H) Representative immunofluorescence images of residual MT staining (DM1A) in WT and Mfn2 KO extracted MEFs. (I) Quantification of residual DM1A tubulin levels in WT and Mfn2 KO MEFs treated as in H (n=100-125 cells). Data are expressed as median with interquartile range. n= 3 independent experiments * p<0.05; ** p<0.01; ns non-significant by Mann–Whitney U test. Scale bars, 10 μm.

We measured MT plus end dynamics by following the behavior of individual MTs in WT and Mfn2 KO cells transfected with GFP-tubulin and found that lack of MFN2 expression almost doubled MT dynamicity, an effect due to an increase in MT growth and shrinkage rates (Table 1). The rise in MT dynamicity correlated with a significant loss of MT stability. To test this, we measured the amount of residual MT polymer resisting depolymerization that was induced by mild detergent extraction prior to fixation and immunofluorescence staining (Fig. 1H,I). Importantly, both acetylated tubulin levels and MT dynamics were normalized in Mfn2 KO cells by the HDAC6 inhibitor trichostatin A (TSA) (Fig. S1A-D and Table S1), suggesting that the increase in MT dynamicity resulted from loss of a-tubulin acetylation in cells deprived of MFN2.

**Table 1.** MFN2 regulates MT dynamics in MEFs. MT dynamics were measured from time-lapse analysis of GFP-tubulin-labeled MTs using epifluorescence microscopy (1f/5s). Parameters characterizing MT dynamics, such as the growth rate, the shrinkage rate, the frequency of ‘catastrophe’ (transitions from growth/pause to shortening) and ‘rescue’ (transitions from shortening to growth/pause) events, as well as the average amount of time spent by MTs in growth, shrinkage and pausing, MT lifetime and MT dynamicity (#of growth + shrinkage events/lifetime). Data are mean ± SEM from 3 independent experiments. * p<0.05; *** p<0.001 by Student’s t-test.

|  | WT | Mfn2 KO |
| --- | --- | --- |
| Growth rate (μm/s) | 0.05 ± 0.02 | 0.13 ± 0.006 * |
| Shrinkage rate (μm/s) | 0.07 ± 0.004 | 0.12 ± 0.01 * |
| Catastrophe freq. (s<sup>-1</sup>) | 0.06 ± 0.006 | 0.06 ± 0.004 |
| Rescue freq. (s<sup>-1</sup>) | 0.08 ± 0.006 | 0.08 ± 0.006 |
| % Growth | 35.5 ± 1.92 | 44.15 ± 0.95 *** |
| % Shrinkage | 34.05 ± 0.87 | 36.48 ± 1.27 * |
| % Pause | 29.58 ± 0.62 | 20.1 ± 1.45 |
| MT lifetime (s) | 58.25 ± 2.14 | 60.5 ± 1.73 |
| MT dynamicity (μm/min) | 5.96 ± 0.42 | 10.43 ± 0.25 *** |
| Number of MTs | 20 | 20 |

### Tubulin acetylation is required for MFN2-dependent regulation of mitochondrial motility but not for mitochondrial fusion or functional tethering to the ER

We observed that the co-localization of mitochondria with MTs was reduced in Mfn2 KO cells but was restored when Mfn2 KO cells were treated with TSA (Fig. S1E,F). Hence, we determined whether increasing acetylated tubulin levels by TSA also reestablished regular mitochondria dynamics, and/or mitochondrial associated ER-membrane (MAM) function, mitochondrial features affected by loss of MFN2 expression (Fig. 2). We observed that TSA normalized both central and peripheral mitochondrial displacement velocity as well as mitochondria distribution and contacts with MTs in Mfn2 KO cells (Fig. 2A-C and Fig. S1E,F). However, while mitochondria elongated morphology was partially reestablished in Mfn2 KO cells treated with TSA, increasing acetylated tubulin completely failed to recover mitochondria fusion, an activity significantly compromised in cells deprived of MFN2 (Fig. 2D,E). Identical results were obtained using tubacin, a more potent and highly selective HDAC6 inhibitor (Fig. 2F-I) [52].

**Figure 2.**
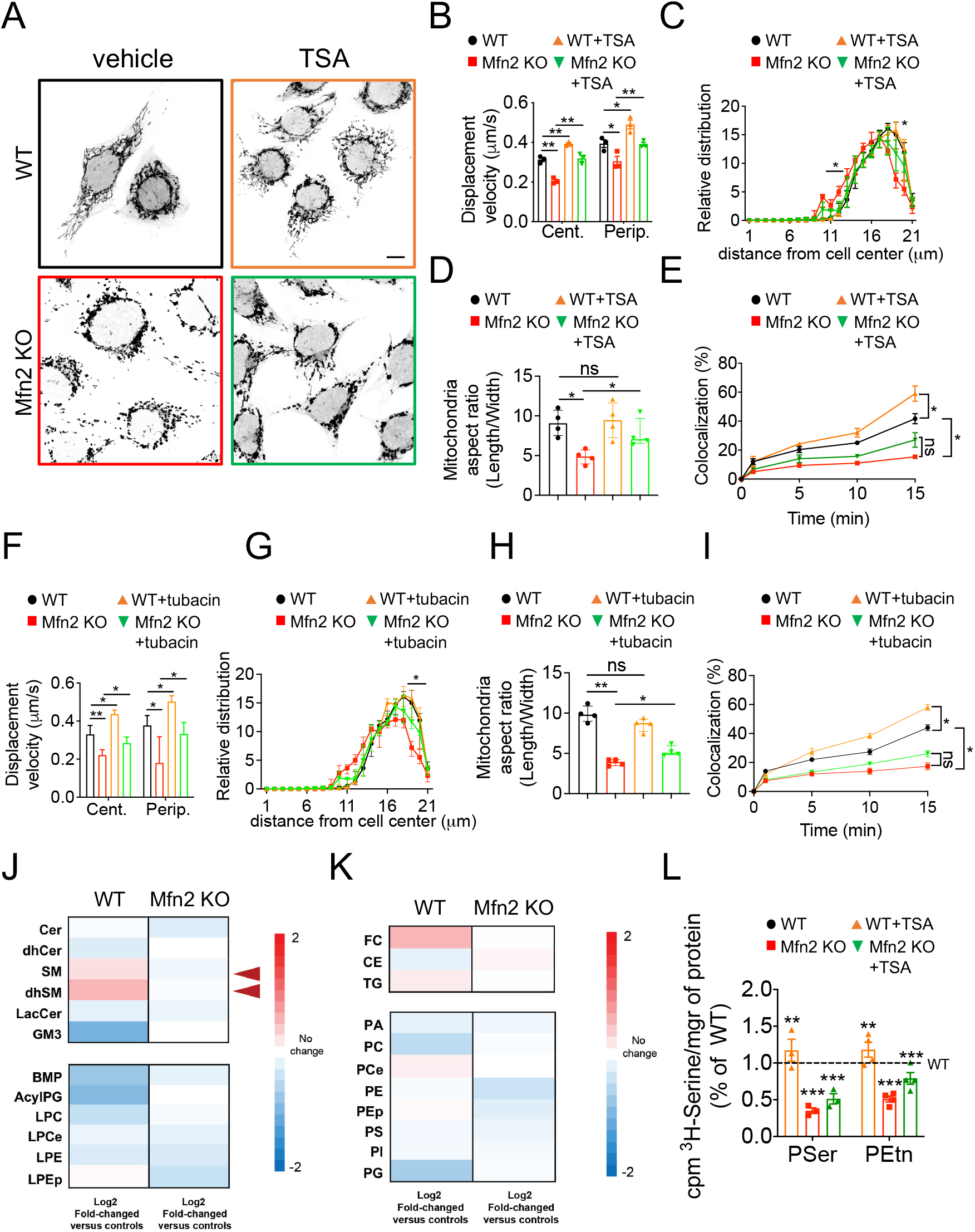
Restoring acetylated tubulin in Mfn2 KO MEFs rescues mitochondria motility and cholesterol esterification but not mitochondria fusion or phospholipid synthesis defects. (A) WT and Mfn2 KO cells were stained with mitoTracker Red and treated with TSA (10 μM) or vehicle control for 6 h. Scale bar, 10 μm. (B) Quantification of mitochondrial displacement velocity analyzed from movies acquired for 3 min (1f/2s) in cells treated as in A. Movies were analyzed using Image J manual tracking plug-in. (C) Quantification of relative distribution of mitochondria (#) to the geometrical cell center in cells treated as in A (D) Quantification of mitochondria aspect ratio (length/width) in cells treated as in A (n = 100-120 mitochondria from 4 independent experiments). (E) Quantification of mitochondria fusion using mitoDendra expression in cells treated as in A. (F) Quantification of mitochondrial displacement velocity analyzed from movies acquired for 3 min (1f/2s) in cells treated with tubacin (20 μM). Movies were analyzed using Image J manual tracking plug-in (125-150 mitochondria/10-12 cells from 3 independent experiments). (G) Quantification of relative distribution of mitochondria (#) to the geometrical cell center in cells treated with tubacin (20 μM). (H) Quantification of mitochondria aspect ratio (length/width) in cells treated with tubacin (20 μM). (I) Quantification of mitochondria fusion using mitoDendra expression in cells treated with tubacin or control vehicle for 6 h. (G,H and I) n= 125-150 mitochondria from 3-4 independent experiments. (J,K) Heatmap representation of changes in lipid classes in MFN2 KO MEFs treated with vehicle control or TSA (10 nM). (L) Phospholipid synthesis and transfer between ER and mitochondria in WT and Mfn2 KO MEFs treated with vehicle control or TSA (10 nM) for 6 h. Incorporation of ^3^H-Ser into ^3^H-PtdSer (PS) and (H) ^3^H-PtdEtn (PE) after 2 h and 4 h expressed as % of the average value measured in the controls. Cer: ceramide, dhcer: dihydroceramide, SM: sphingomyelin, dhSM: dehydrosphingomyelin, GM3: monosialodihexosylganglioside. BMP: Bis(monoacylglycerol)phosphate, Acyl-PG, acylated phosphatidylglycerol, LPC: Lysophosphatidylcholine, LPCe: Lysophosphatidylcholine plasmalogen, LPE: Lysophosphatidylethanolamine, LPEp: Lysophosphatidylethanolamine plasmalogen; FC: free cholesterol, CE: cholesteryl esters; PA: phosphatidic acid; PC: Phosphatidylcholine; PCe Phosphatidylcholine plasmalogen; PE: Phosphatidylethanolamine; PEp: Phosphatidylethanolamine plasmalogen; PS: Phosphatidylserine; PI: Phosphatidylinositol; PG: Phosphatidylglycerol. * p<0.05; ** p<0.01; *** p<0.001; ns non-significant by Kruskal-Wallis test. Data are expressed as median with interquartile range.

We inquired whether the rescue of mitochondrial dynamics was dependent on acetylated tubulin or a general gain in MT stability resulting from tubulin acetylation. To test this, we adopted Iqgap1 KO MEFs, a cell line with normal MFN2 levels but naturally deprived of detyrosinated and acetylated MTs, two independent subsets of stable MTs [53] (Fig. S2). By analogy with Mfn2 KO cells, we found that loss of IQGAP1 also resulted in defective MT and mitochondrial dynamics (Fig. S3 and Table S2), consistent with a role for modified MTs in regulating mitochondria homeostasis. However, increasing detyrosinated tubulin by tubulin tyrosine ligase (TTL) silencing did not normalize mitochondrial dynamics to the extent of TSA treatment (Fig. S3E-I), suggesting that rescue of mitochondria dynamics was dependent on the selective increase in acetylated tubulin rather than a general gain in MT stability (Fig. S2,3 and Table S2).

Next, we tested the effects of restoring acetylated tubulin levels on loss of MAM function by analyzing the synthesis and transfer of phospholipid between ER and mitochondria, a known proxy of MAM activity, as well as changes in lipid classes by lipidomics analysis in Mfn2 KO cells [34, 54]. MAM is a transient specialized subdomain of the ER with the characteristics of a lipid raft. The temporary formation of MAM domains in the ER regulates several metabolic pathways, including lipid and Ca^2+^ homeostasis and mitochondrial activity [38, 55]. Alterations in the formation of MAM domains have been reported to induce significant changes in lipid metabolism in several pathologies including neurodegenerative disease [39, 55]. In particular, defects in MAM activity have significant detrimental effects on the regulation of cholesterol and its esterification into cholesteryl esters [56]. Equally important, defects in MAM impair the regulation of sphingomyelin (SM) turnover and its hydrolysis into ceramide species [57, 58].

We found that Mfn2KO cells display significant increases in sphingomyelin and cholesterol with concomitant decreases in cholesteryl esters and ceramide levels (Fig. 2J,K) and that these changes could be rescued by HDAC6 inhibition (Fig. 2J,K). However, TSA failed to normalize MAM-dependent phospholipid synthesis measured by incorporation of radiolabeled ^3^H-Ser into newly synthesized ^3^H-PtdSer (PS) and (H) ^3^H-PtdEtn (PE) (Fig. 2L).

Altogether, our results demonstrate a previously unrecognized role for MFN2 in the regulation of α-tubulin acetylation and suggest that this activity is important for MFN2-dependent control of mitochondria motility and lipid-raft MAM composition, but not for MFN2-dependent mitochondrial fusion or functional mitochondrial/ER tethering. Furthermore, our results in Iqgap1 KO cells support the notion that acetylated tubulin is a modulator of mitochondria dynamics *per se* and suggest that the machinery controlling mitochondria motility may regulate the α-tubulin acetylation cycle at sites of mitochondria contacts with MTs.

### MFN2 regulates α-tubulin acetylation by recruiting ATAT1 at sites of mitochondrial contacts with MTs

We began to investigate the mechanisms underlying MFN2 regulation of acetylated α-tubulin by measuring levels and localization of ATAT1 and HDAC6 in Mfn2 KO cells. HDAC6 expression was three-fold higher in these cells, in contrast to ATAT1 levels, which remained unaffected (Fig. S4A,B). Loss of MFN2 expression did not affect the percentage of cells in mitosis either (Fig. S4C,D). However, when intracellular membranes were subjected to crude fractionation to isolate the cytosolic from the nuclear and ER fractions, unlike HDAC6 which remained mostly cytosolic, ATAT1 appeared in the cytosolic and in the nuclear/ER portion in WT cells but re-distributed more prominently to the nuclear/ER fraction in Mfn2 KO cells (Fig. S4E-G). Accordingly, co-localization of endogenous ATAT1 with the ER appeared to be increased in Mfn2 KO cells compared to WT cells (Fig. S4H-J).

We hypothesized that MFN2 may negatively regulate ATAT1 association with the ER by localizing ATAT1 to mitochondria outer membranes, and that this localization may facilitate the access of ATAT1 to openings of the MT lattice at sites of mitochondria contacts with MTs. High resolution confocal microscopy of endogenous proteins revealed punctuate localization of ATAT1 to mitochondria membranes or MFN2, and this co-localization was lost in cells deprived of MFN2 expression (Fig. 3A-F). Localization of ATAT1 to mitochondria was likely to be dependent on the association of MFN2 with an ATAT1 N-terminal fragment (1-242) inclusive of its catalytic domain, as demonstrated by the *in situ* validation of this interaction using the proximity ligation assay (Fig. 3G,H) and conventional pull down analyses from whole cell lysates using full length or C-terminally truncated versions of ATAT1 (Fig. 3I-K).

**Figure 3.**
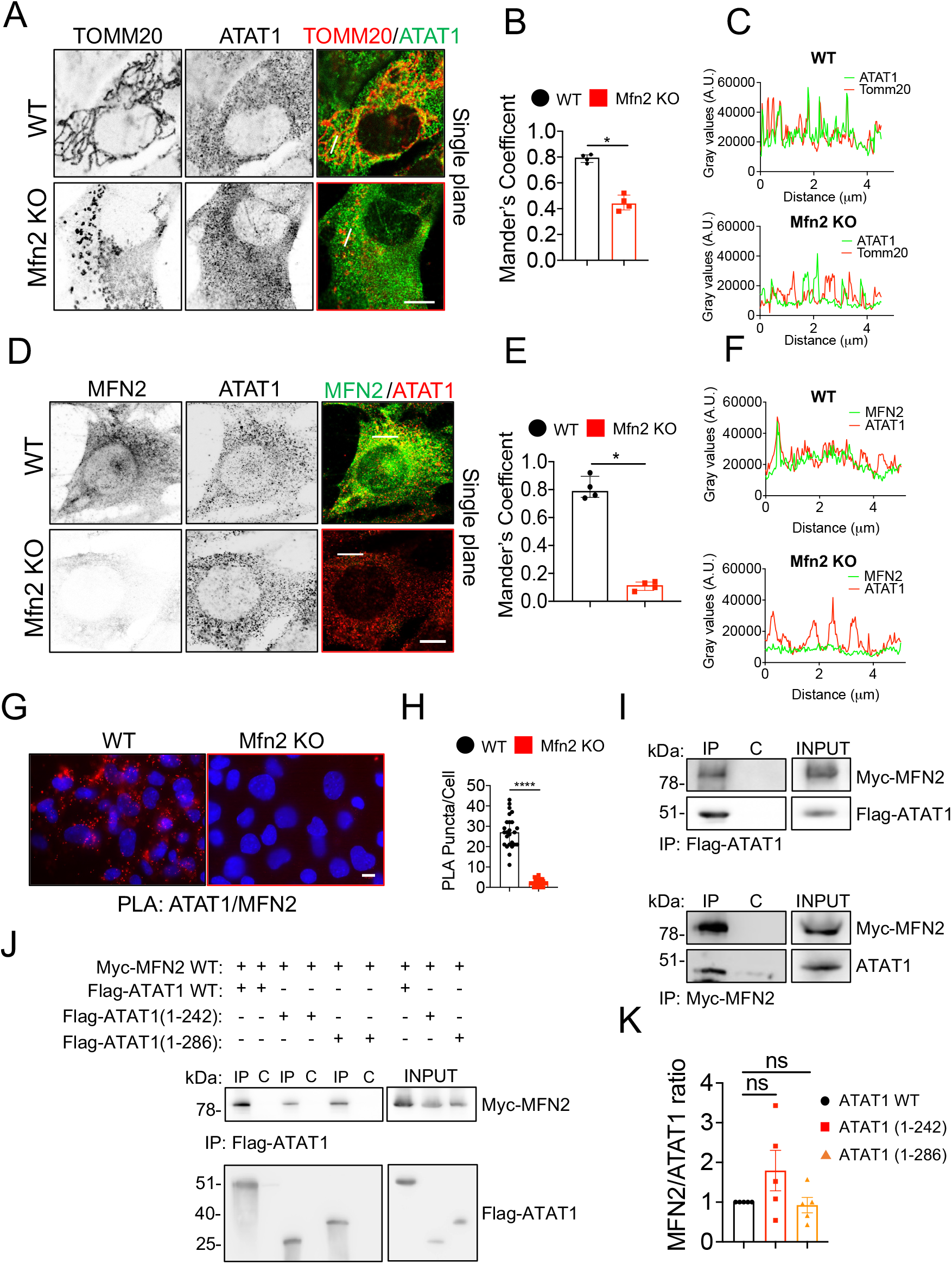
MFN2 localizes the ATAT1 to mitochondria outer membranes in MEFs. (A) Airyscan confocal analysis of mitochondria (TOMM20) and ATAT1 in WT and Mfn2 KO MEFs. Scale bar, 5 μm. (B) Quantification of localization of ATAT1 at mitochondria as in (A) by Mander’s correlation coefficient. (C) Line scan analysis of mitochondria and ATAT1 localization from selected regions in (A). Lines are shown as white bars in (A). (D) Airyscan confocal analysis of ATAT1 and MFN2 localization in WT and Mfn2 KO MEFs. Scale bar, 5 μm. (E) Quantification of localization of ATAT1 and MFN2 as in (D) by Mander’s correlation coefficient. (F) Line scan analysis of MFN2 and ATAT1 localization from selected regions in (D). Lines are shown as white bars in (D). (G) Immunofluorescence analysis of MFN2 and ATAT1 PLA signal in WT and Mfn2 KO MEFs. Scale bar, 10 μm. (H) Quantification of PLA puncta per cell in WT and Mfn2 KO MEFs. n= 50-55 cells from 3 independent experiments. (I) Interaction between ATAT1 and MFN2 was detected by immunoprecipitation (IP) followed by immunoblot analysis with the indicated antibodies (upper panel). Similarly, interaction between endogenous ATAT1 and transfected Myc-MFN2 WT was detected in HEK293T cells (lower panel). (J) HEK293T cells were co-transfected with Myc-MFN2 WT and Flag-ATAT1 WT or Flag-ATAT1 (1-242) or Flag-ATAT1 (1-286). Interaction between MFN2 and ATAT1 WT or mutants was detected by co-immunoprecipitation followed by immunoblot analysis with the indicated antibodies. (K) Normalized MFN2 to ATAT1 signal ratio from (J) was plotted. Data are expressed as median with interquartile range. n = 35-50 cells from 3-4 independent experiments. * p<0.05; **** p<0.001; ns non-significant by Mann–Whitney U test (3B,E and H) and Kruskal-Wallis test (3K).

Altogether, our data demonstrate that ATAT1 associates with mitochondria and that this localization is dependent on the binding of the catalytic domain of ATAT1 with MFN2.

### Loss of acetylated tubulin may underlie CMT2A disease

Most MFN2 CMT mutations are missense, and all produce a dominant inheritance pattern, suggesting that mutations in MFN2 lead to either a gain of function or haploinsufficiency [59, 60]. Furthermore, recent work supports the notion that restoring MNF1:MFN2 balance by increasing levels of its homologous protein MFN1 is a potential therapeutic approach for CMT2A [46]. The reason for this compensation is unclear, although both MFN2 and MFN1 have been implicated in mitochondria fusion. To determine the involvement of MFN2-dependent regulation of tubulin acetylation in CMT2A disease we investigated whether: 1) mutations in MFN2 affect the interaction with ATAT1 and/or fail to restore normal acetylated tubulin levels in Mfn2 KO cells; 2) MFN1 compensates for loss of MFN2 by restoring tubulin acetylation in Mfn2 KO cells; 3) loss of acetylated tubulin by MFN2 depletion is conserved in sensory neurons and sufficient to induce axonal fragmentation, a phenotype associated to axonal forms of CMT disease including CMT2A.

We found that MFN2 R94W and T105M, two of the most common N-terminal CMT mutations in MFN2 [61–63] bind to ATAT1 with higher affinity than WT MFN2 (Fig. 4A-C). In addition, while immunofluorescence analysis showed significant higher co-localization of endogenous ATAT1 only with the transfected T105M mutant (Fig. 4D,E), both mutations failed to rescue normal acetylated tubulin levels when expressed in Mfn2 KO cells (Fig. 4F,G). This result was in contrast with ectopic expression of MFN1, which was able to compensate for loss of MFN2 on acetylated tubulin levels in Mfn2 KO cells (Fig. 4H,I).

**Figure 4.**
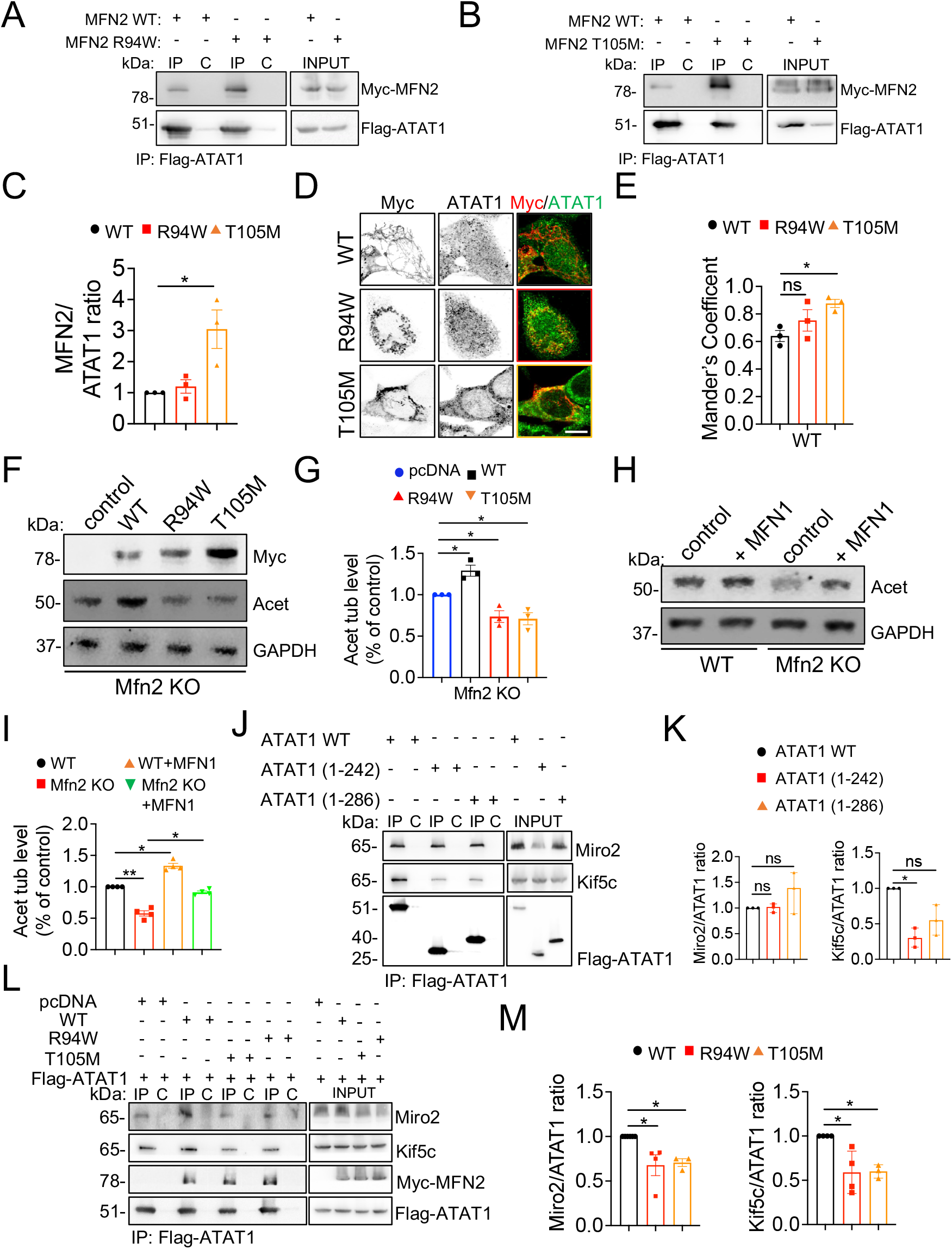
Regulation of a-tubulin acetylation by MFN2 is affected by MFN2 mutations and shared by MFN1. (A) HEK293T cells were co-transfected with Flag-ATAT1 and Myc-MFN2 WT or Myc-MFN2 R94W. (B) HEK293T cells were co-transfected with Flag-ATAT1 and Myc-MFN2 WT or Myc-MFN2 T105M. Interaction between ATAT1 WT and MFN2 WT or mutants was detected by immunoprecipitation (IP) followed by immunoblot (IB) with the indicated antibodies. (C) The ratio of the MFN2 signal to the ATAT1 signal from (A and B) was plotted. (D) Immunofluorescence analysis of overexpression of Myc-MFN2 WT, Myc-MFN2 R94W, Myc-MFN2 T105M in WT MEF cells (n=25-30 cells). (E) Mander’s coefficient analysis for WT MEF cells co transfected with Myc-MFN2 WT, Myc-MFN2 R94W and Myc-MFN2 T105M. (F) representative immunoblot of acetylated tubulin levels in Mfn2 KO cells co-transfected with Myc-MFN2 WT, Myc-MFN2 R94W and Myc-MFN2 T105M. (G) Quantification of acetylated tubulin levels expressed as % of control levels from 3 independent experiments as in F. (H) Representative immunoblot of acetylated tubulin levels in MFN1 overexpressing WT and Mfn2 KO cells. (I) Quantification of acetylated tubulin levels expressed as % of control levels from 4 independent experiments as in H. (J) Interaction between ATAT1 or its truncated mutants (1-242; 1-286) and endogenous Miro2 or Kif5c was detected in HEK293T cells transfected with Flag-ATAT1 WT or its truncated mutants. (K) The ratio of the Miro2 and Kif5c signal to the ATAT1 signal from (J) was plotted. (L) Interaction between Flag-ATA1 and endogenous Miro2 or Kif5c was detected in HEK293T cells in presence or absence of Myc-MFN2 WT, Myc-MFN2 R94W and Myc-MFN2 T105M. (M) The ratio of the Miro2 and Kif5c signal to the ATAT1 signal from (L) is plotted. Data are expressed as median with interquartile range. n= 3-4 independent experiments. * p<0.05, ** p<0.01; ns non-significant by Kruskal-Wallis test. Scale bar, 10 μm.

A complex between miro/Milton (TRAK) and MFN2 has been previously shown, and miro has been implicated in regulating MFN2-dependent mitochondrial fusion in response to mitochondrial Ca^2+^ concentration [43]. We tested whether also ATAT1 interacted with miro and/or kinesin heavy chain (Kif5c) and determined the potential effects of mutant MFN2 on the formation of these complexes. We found that ectopic ATAT1 co-immunoprecipitated with both endogenous miro2 and kif5c and that ectopic expression of mutant MFN2 R94W or T105M significantly lowered the affinity of these bindings (Fig. 4J-M). Taken together, these data demonstrate that regulation of acetylated tubulin is an activity shared by MFN1 and that loss of acetylated tubulin may play a primary role in CMT2A *via* the sequestering effect of MFN2 mutations on ATAT1 binding.

These observations became particularly meaningful when we tested the consequences of loss of MFN2 in sensory neurons and the effects of HDAC6 inhibition on these phenotypes. By analogy with Mfn2 KO cells, silencing of MFN2 expression reduced acetylated tubulin levels both in adult mouse DRG neurons grown in culture and in cell bodies of somatosensory neurons of third instar stage *Drosophila* larvae (Fig. 5A-D). Similar to Mfn2 KO MEFs, we observed localization of endogenous ATAT1 and MFN2 in DRG neurons (Fig. S5A-C) and significant reduction in the extent of ATAT1 localization to mitochondrial membranes in neurons silenced for Mfn2 expression in both proximal and distal portion of the axon (Fig. S5D-F). Importantly, cultured sensory neurons deprived of MFN2 acquired a dying-back degeneration phenotype starting from distal regions of the axon, as indicated by the appearance of retraction bulbs at the onset of axonal fragmentation (Fig. 5E and F). We observed that loss of acetylated tubulin preceded axonal degeneration in DRG neurons deprived of MFN2 for shorter times (Fig. S5G-J) and that HDAC6 inhibition, which significantly rescued normal acetylated tubulin levels in Mfn2 KD neurons, prevented both retraction bulb formation and axonal degeneration in neurons deprived of MFN2, while having only negligible effects on WT controls (Fig. 5G-I).

**Figure 5.**
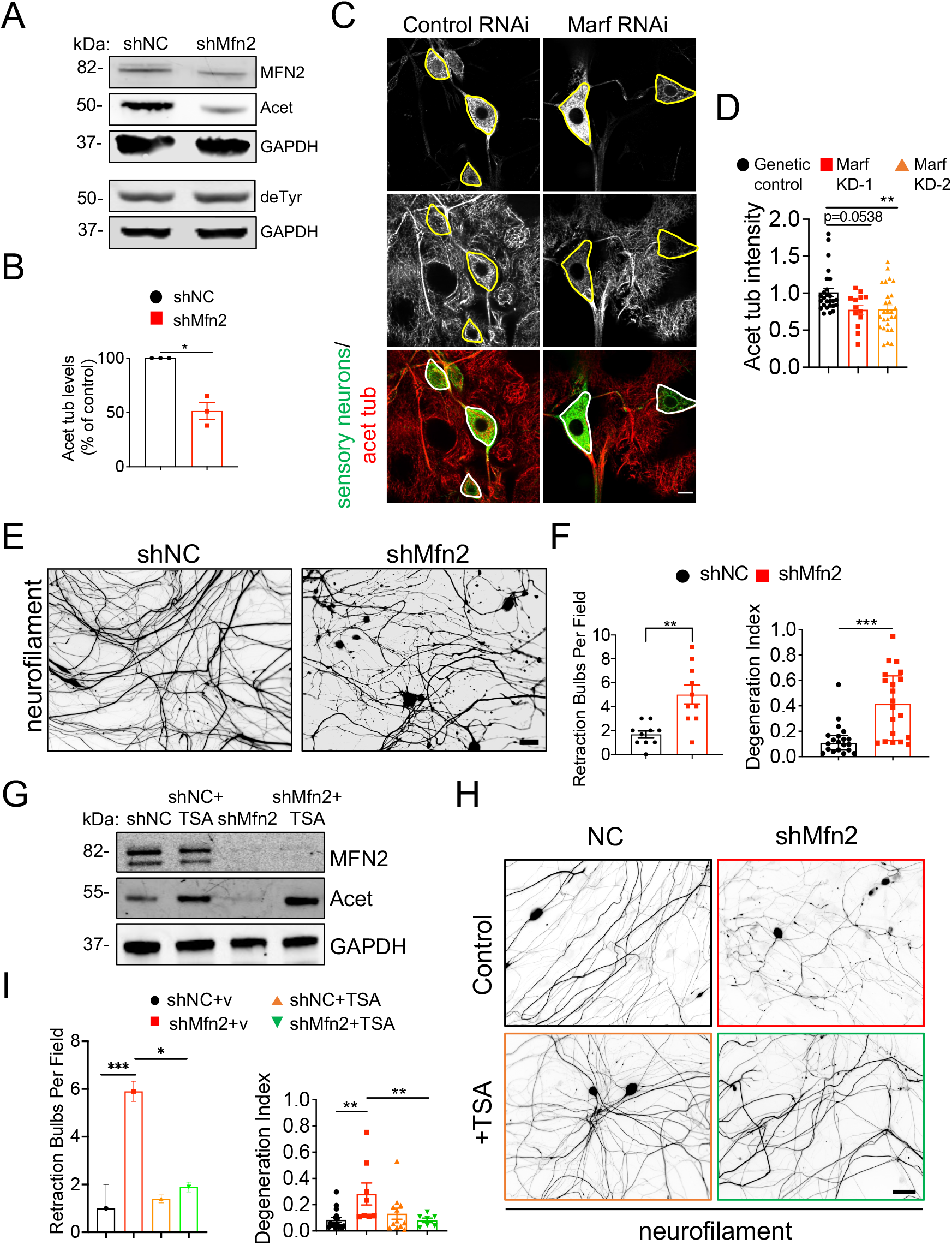
MFN2 regulates α-tubulin acetylation in sensory neurons in vitro and in vivo and this activity is required for axonal integrity. (A) Representative immunoblot of MFN2, acetylated (Acet) and detyrosinated tubulin (deTyr) levels in adult DRG neurons (14 DIV) silenced of Mfn2 expression at 7 DIV. GAPDH, loading control. (B) Quantification of acetylated tubulin relative in MFN2 depleted DRG neurons relative to control neurons infected with shNC (non-coding shRNA). (C) Multi-dendritic neuron driver 109(2)80 Gal4 driver was used to label somatosensory neurons (CD8-GFP) and knockdown (KD) Marf in *Drosophila* larval somatosensory neurons. Acetylated tubulin levels were measured in the cell bodies as demarcated in the images. Confocal stacks spanning the cell body were examined to reveal acetylated tubulin staining in cell bodies distinct from nearby epidermis. To quantify levels of acetylated tubulin staining in cell bodies, images were sub-stacked for each cell body (average intensity z-projection) and blinded. Mean gray value was measured and normalized against the background level of acetylated tubulin staining (see methods for further information). (D) Quantification of acetylated tubulin protein expression in cell bodies after Marf knockdown in somatosensory neurons in larvae. Two different strategies were used, Marf KD-1 (BL 67158) and KD-2 (BL 31157). (E) Images of representative fields showing dissociated adult DRG neurons (14 DIV) treated as in A, fixed and immunostained with mouse anti-neurofilament (2H3-s) antibody. (F) Quantification of number of retraction bulbs per field and degree of axonal degeneration. The area occupied by the axons (total axonal area) and degenerating axons (fragmented axonal area) was measured in the same field from images in WT and MFN2 KD DRG neurons. Degeneration index was calculated as the ratio between fragmented axonal area and total axonal area. (G) Representative immunoblot of MFN2, acetylated tubulin (Acet) levels in control (shNC) and MFN2 (shMFN2) silenced DRG neurons incubated with 10 μM of the HDAC6 inhibitor TSA or vehicle control for 6 h prior to lysis. GAPDH, loading control. (H) Representative immunofluorescence images of DRG neurons treated as in G. (I) Quantification of number of retraction bulbs per field and degree of axonal degeneration in DRG neurons treated as in G prior to fixation and staining. Data are represented as median and interquartile range from 3 independent experiments. * p<0.05; ** p<0.01, *** p<0.001 by Mann-Whitney U test (B, D and F) and Kruskal-Wallis test (I). Scale bars, 5 μm (C); 50 μm (H).

These findings indicated that MFN2 dependent recruitment of ATAT1 to sites of mitochondrial contacts with MTs is conserved in sensory neurons and required for axonal integrity by maintaining normal levels of MT acetylation. Taking consideration of our functional data in MEF cells, and consistent with previous observations in cellular models of CMT2 caused by MFN2 mutations, these results also suggest that distal axonal degeneration caused by mutant MFN2 predominantly depends on loss of acetylated tubulin, which affects mitochondrial motility and distribution, but not on loss of fusion or functional mitochondria/ER tethering.

## Discussion

MFN2 mutations in CMT2A disrupt the fusion [64] of mitochondria and compromise ER-mitochondrial interactions [34, 47]. However, while certain CMT2A mutant forms of MFN2 impair mitochondrial fusion and/or functional mitochondria/ER tethering, others do not affect either function [64], casting doubt on the implication of these MFN2 activities in the etiology of CMT2.

In this study we report that MFN2 is a regulator of a-tubulin acetylation and MT dynamics, and that in Mfn2 KO MEFs rescuing α-tubulin acetylation levels by pharmacological inhibition of HDAC6 corrects defects in MT dynamics and mitochondrial motility, some MAM function but not MAM integrity or mitochondrial fusion. We also show that regulation of tubulin acetylation by MFN2 occurs through MFN2-mediated recruitment of ATAT1 to outer mitochondrial membranes, an activity conserved in sensory neurons, critical in the induction of axonal degeneration by MFN2 loss of function and impaired in two MFN2 mutants associated with CMT2A. Interestingly, the binding of MFN2 to ATAT1 is dependent on the N-terminal catalytic domain of ATAT1 and the same domain is also necessary for the association of ATAT1 with kinesin-1 but not with miro, a Rho-GTPase implicated in the regulation of mitochondrial transport by linking mitochondria outer membranes to kinesin and dynein motors [65, 66]. Conversely, both MFN2 R94W and T105M mutants disrupt the binding of ATAT1 with either miro or kinesin-1, suggesting that while ATAT1 binding to miro may not depend on kinesin, the formation of a stable ATAT1/miro/kinesin-1 complex relies on functional MFN2. Based on these observations, we propose that, in analogy to axonal vesicles [21], mitochondria contacts with MTs are hotspots of tubulin acetylation and that this function is impaired in CMT2 disease caused by MFN2 mutations. Specifically, we suggest that mutant MFN2 R94W or T105M drive axonal degeneration by disrupting the ability of mitochondria to release the ATAT1 at specific sites on axonal MTs, leading to an imbalance in tubulin acetylation and disrupted mitochondrial transport. Our findings also provide evidence that the release of ATAT1 by MFN2 depends on the formation of a stable ATAT1/miro/kinesin-1 complex, which may be necessary to allow discharge of ATAT1 at putative entry sites into the MT lattice. This is in line with the observation that motors can leave marks in the MT shaft by inducing breaks in the lattice and promote MT self-repair [48, 50, 51]. Further work is required to understand the rules of site selection and whether a break in the MT lattice is sufficient to induce ATAT1 release from MFN2 through a putative conformational change in the motor complex.

By combining our observations in Mfn2 KO MEFs and KD sensory neurons, we propose that axonal degeneration caused by MFN2 loss of function mutations may not depend on impaired mitochondrial fusion or functional mitochondria/ER tethering, but rather loss of MFN2-dependent regulation of mitochondrial transport by interfering with tubulin acetylation at sites of mitochondria and MT contact. Our interpretation is consistent with a pathogenic role for disrupted mitochondrial transport in neuropathies and a key role for tubulin acetylation in mitochondrial dynamics. Indeed, multiple studies report that axonal degeneration precedes cell body death in several peripheral neuropathies, including CMT disease. Mitochondria are the principal mediators of ATP production and Ca^2+^ buffering, and they actively distribute to areas of high energy demand and Ca^2+^ flux within the axon [43, 44]. A general defect in the ability of mitochondria to translocate to these sites would be expected to lead to preferential degeneration of long axons that frequently experience fluctuations in ATP and Ca^2+^ levels.

Several lines of evidence further support a role for perturbation of acetylated tubulin levels in CMT2A: 1) mutant MFN2 (MFN2R94Q) knock-in mice lack acetylated tubulin in distal axons of their long peripheral nerves [29]; 2) HDAC6 inhibition has been reported to be a promising therapeutical approach in several toxic and familial peripheral neuropathies, including CMT2A [29]; 3) the formin INF2, mutations of which cause dominant intermediate CMT in association with FSGS [67], is a positive regulator of tubulin acetylation by modulating ATAT1 transcription. We note, however, that HDAC6 deacetylates additional lysine residues of α- and β-tubulin and has multiple substrates in addition to tubulin, casting doubt on the specificity of this approach. Conversely, tubulin is the only known substrate for ATAT1. In addition, the structure of ATAT1 provides a unique scaffold for designing small molecule modulators of tubulin acetylation for therapeutic use [68]. In particular, the identification of ATAT1 mutations that decrease or increase ATAT1 activity suggests that small molecule compounds could be identified to increase or decrease ATAT1 activity to stabilize or destabilize MTs for therapeutic purposes [68]. Taken together, these studies indicate that targeting ATAT1 activity or expression may represent an alternative and more specific therapeutic approach aimed at restoring sensory neuron function in CMT2A and perhaps other related CMT subtypes.

## MATERIALS AND METHODS

| REAGENT or RESOURCE | SOURCE | IDENTIFIER |
| --- | --- | --- |
| **Antibodies** |  |  |
| Mouse monoclonal mitofusin-2 (6A8) | Abcam | Ab56889 |
| Rabbit monoclonal mitofusin-2 | Abcam | Ab124773 |
| Rabbit polyclonal IQGAP1 | Novus<br>Biologicals | NBP1-06529 |
| Rabbit polyclonal HDAC6 | Novus<br>Biologicals | NBP1-78981 |
| Rabbit polyclonal ATAT1 | Bioss | Bs-9535R |
| Rabbit monoclonal TOMM20 | Abcam | Ab186735 |
| Mouse monoclonal TOMM20 | Millipore Sigma | KB061-4F3 |
| Rabbit polyclonal detyrosinated tubulin | Abcam | Ab48389 |
| Rat tyrosinated tubulin, clone YL1/2 | Millipore | MAB1864-I |
| Mouse monoclonal acetylated tubulin | Sigma Aldrich | T6793 |
| Mouse monoclonal DM1A | Sigma Aldrich | T6199 |
| Human anti-tubulin GTP (MB11) | Adipogen | AG-27B-0009-C100 |
| Rabbit polyclonal TTL | Proteintech | 13618-1-AP |
| Mouse monoclonal PDI | Santacruz | SC-74551 |
| Mouse monoclonal anti-c-Myc 9E10 HRP | Santacruz | SC-40 HRP |
| Mouse monoclonal anti-c-Myc 9E10 Agarose | Santacruz | SC-40 AC |
| Mouse monoclonal anti-Flag M2 HRP | Sigma Aldrich | A8592 |
| Mouse monoclonal anti-Flag M2 Agarose | Sigma Aldrich | A2220 |
| Chicken polyclonal neurofilament | Aves labs | AB_2313553 |
| Rabbit polyclonal anti-Miro2 | Proteintech | 11235-1-AP |
| Rabbit polyclonal anti-Kif5c | Proteintech | 25897-1-AP |
| Mouse monoclonal GAPDH | Abcam | Ab8245 |
| Rabbit polyclonal GAPDH | ThermoFisher | PA1-987 |
| Goat anti-mouse IgG (H+L) highly cross-adsorbed secondary antibody, Alexa Flour 488-conjugated | ThermoFisher | A11029 |
| Goat anti-rabbit IgG (H+L) highly cross-adsorbed secondary antibody, Alexa Flour 488-conjugated | ThermoFisher | A11034 |
| Goat anti-mouse IgG (H+L) highly cross-adsorbed secondary antibody, Alexa Flour 546-conjugated | ThermoFisher | A11030 |
| Goat anti-rabbit IgG (H+L) highly cross-adsorbed secondary antibody, Alexa Flour 546-conjugated | ThermoFisher | A11035 |
| Goat anti-human IgG (H+L) highly cross-adsorbed secondary antibody, Alexa Flour 568-conjugated | ThermoFisher | A-21090 |
| IRDye® 680RD Goat anti-mouse IgG secondary antibody | LI-COR | 926-68070 |
| IRDye® 800CW Goat anti-rabbit IgG secondary antibody | LI-COR | 926-32211 |
| IRDye® 680RD Goat anti-rabbit IgG secondary antibody | LI-COR | 926-68071 |
| IRDye® 800CW Goat anti-rabbit IgG secondary antibody | LI-COR | 926-32211 |
| **Bacteria strains** |  |  |
| DH5 alpha | New England Biolabs | C2987I |
| XL1-Blue | Agilent | 200229 |
| **Chemicals and Reagents** |  |  |
| Trichostatin A (TSA) | Tocris | #1406 |
| Tubacin | Millipore Sigma | SML0065 |
| Mitotracker Red CMXROS | Thermofisher | M7512 |
| DMEM | Gibco | 11995-065 |
| Neurobasal | Thermofisher | 110349 |
| Fetal bovine serum | HyClone | SH30071.03 |
| Bovine calf serum | Thermofisher | 26170043 |
| B-27 supplement (50x) | Thermofisher | 7504044 |
| Penicillin-streptomycin | Thermofisher | 15140163 |
| 10x HBSS | Thermofisher | 14065056 |
| Cytosine β-D-arabinofuranoside hydrochloride (AraC) | Millipore Sigma | C6645 |
| Collagenase | Millipore Sigma | C0130 |
| Trypsin 0.05% EDTA | Thermofisher | 25300054 |
| GlutaMAX Supplement | Thermofisher | 35050061 |
| Poly-D-Lysine | Sigma Aldrich | P1149 |
| Laminin | Sigma Aldrich | 11243217001 |
| Laemlli SDS sample buffer, reducing | Thermofisher | J60015-AD |
| NuPAGE MOPS SDS Running buffer | Thermofisher | NP0001 |
| Fluoromount-G | Southern Biotech | 0100-01 |
| 32% PFA | EMS | 15714-S |
| DreamFect™Gold transfection reagent | OZ Bioscience | DG80500 |
| c-Myc Peptide | Sigma Aldrich | M2435 |
| Flag peptide | Sigma Aldrich | F3290 |
| **Critical commercial assays** |  |  |
| DuoLink PLA In Situ Red starter mouse/rabbit kit | Sigma-Aldrich | DUO92101 |
| **Experimental models: Organisms/strains** |  |  |
| *UAS-MARF RNAi* | Bloomington<br>Stock Center | BL 31157 |
| *UAS-MARF miRNA* CDS | Bloomington<br>Stock Center | BL 67158 |
| Mouse: C57BL/6J | Charles River<br>Laboratories | RRID:IMSR_CRL:<br>027 |
| **Recombinant DNA** |  |  |
| pLKO.1 shMfn2 | Sigma Aldrich | TRCN0000080608 |
| pLKO.1 shTTL | Sigma Aldrich | TRCN0000191515 |
| Myc-MFN2 WT | David Chan | NA |
| Myc-MFN2 R94W | David Chan | NA |
| Myc-MFN2 T105M | David Chan | NA |
| Flag-ATAT1 WT | Laurent Ngyuen | NA |
| Flag-ATAT1 (1-242) | Laurent Ngyuen | NA |
| Flag-ATAT1 (1-286) | Laurent Ngyuen | NA |
| Zsgreen-MFN1 | Estella Area<br>Gomez | NA |
| **Software** |  |  |
| ImageJ (Fiji) | NIH | RRID:SCR_00228<br>5 |
| GraphPad Prism | GraphPad | RRID:SCR_002798 |
| Li-COR Image Studio Software | Li-COR | RRID:SCR_015795 |
| Zeiss ZEN | Zeiss | RRID:SCR_013672 |
| Andor iQ3 | Oxford Instruments | RRID:SCR_014461 |
| **Other** |  |  |
| 18 mm No.1 circle coverglass | Carolina | 633033 |
| 35 mm MatTek dishes | MatTek | P35G-1.5-14-C |
| NuPAGE Gel | ThermoFisher | NP0316 |
| Nitrocellulose membrane | Fisher | 10600011 |

### Lead contact and materials availability

Further information and requests for resources and reagents should be directed to and will be fulfilled by the lead contact, Francesca Bartolini.

### Experimental model and subject details

All protocols and procedures for mice were approved by the Committee on the Ethics of Animal Experiments of Columbia University and according to Guide for the Care and Use of Laboratory Animals of the <u>National Institutes of Health</u>.

### Cell culture and analyses

WT, Mfn2 KO, Mfn1 KO (kind gifts of Dr. Area-Gomez) and Iqgap1 KO mouse embryonic fibroblast cells were grown in DMEM supplemented with 10% fetal bovine serum. Cells were grown to 80% confluency on acid treated glass coverslips prior to experiment.

### Immunofluorescence microscopy and analyses

For immunofluorescence staining of the MT cytoskeleton, cells were fixed in ice cold MetOH for 10’ prior to rehydration in PBS buffer o/n at 4°C. For all other stainings, cells were fixed in 4% PFA for 15 min and permeabilized with 0.1% Triton X-100 for 5 min at R.T. Cells were then washed in PBS, blocked in 2% FBS and 2% BSA in PBS for 1 h, stained with primary antibodies overnight at 4°C followed by secondary antibodies for 1 h. Mounted samples were observed using a Zeiss LSM 800 confocal microscope equipped with Airyscan module, using a 63× objective (Plan-Apochromat, NA 1.4). Images were acquired and processed using Zen Blue 2.1 software. All images were analyzed by ImageJ software.

### Western blot analyses

Cells were lysed in Laemmli sample buffer and boiled at 96°C for 5 min. Cell lysates were sonicated with a probe sonicator to sheer cellular debris and genomic DNA. Proteins were separated by 10% Bis-Tris gel (Invitrogen) and transferred onto nitrocellulose membrane. After blocking in 5% milk/TBS or BSA/TBS, membranes were incubated with primary antibodies at 4°C overnight prior to 1 h incubation with secondary antibodies. Image acquisition was performed with an Odyssey imaging system (LI-COR Biosciences, NE) and analyzed with Odyssey software.

### Proximity ligation assay (PLA)

PLA assays were carried out using a Duolink *in situ* red starter kit mouse/rabbit kit (Sigma-Aldrich) according to the manufacturer’s protocol. The primary antibodies used were mouse anti-MFN2 and rabbit anti-ATAT1 (1:1000 dilution). Images were acquired on a Zeiss LSM 800 confocal microscope and analyzed using ImageJ/FIJI. Data were pulled from at least three independent biological repeats.

### Analysis of mitochondrial morphology, motility and distribution

Mitochondria were labeled using mitotracker Red CMROX according to manufacturer protocol (Thermo Fisher Scientific) and detected by epifluorescence microscope equipped with 60 x objective lens (Olympus IX81) and a monochrome CCD camera (Sensicam QE, Cooke Corporation). Aspect ratio (length/width) were measured using Image J/FIJI. For mitochondrial distribution and displacement velocity mitochondria were live imaged for 3 min at 2 sec/frame at 37°C. A customized Mitoplot software (kind gift of Dr. Gregg Gundersen) was used for analyzing mitochondrial distribution. Manual tracking plug-in in ImageJ/FIJI was used to analyze mitochondrial displacement velocity.

### Microtubule dynamics

Fibroblasts were transfected with pMSCV-puro-tagGFP-C4 α-tubulin plasmid to generate a green fluorescent protein (GFP)–tubulin stably expressing cell line. Live imaging of MT dynamics in transfected cells was performed at 37°C and 5% CO_2_ for 5 min (5 s/frame) with a 100× PlanApo objective (numerical aperture 1.45) and an iXon X3 CCD camera (Andor, Belfast, United Kingdom) on a Nikon Eclipse Ti microscope controlled by Nikon’s NIS-Elements software (Nikon, Tokyo, Japan). Movies were analyzed by ImageJ using a manual tracking plug-in.

### Mitodendra

Dendra2 photoconversion and imaging utilized the protocol from Evrogen. Images were acquired with an Olympus spinning disk microscope EC-Plan-Neofluar 40X/1.3 oil. Z-stack acquisitions over-sampled each optical slice twice, and the Zen 2009 image analysis software was used for maximum z-projections. The 488 nm laser line and the 561 nm laser excited Dendra2 in the unconverted state and photo-converted state, respectively. To photo-switch Dendra2, a region was illuminated with the 405 nm line (4% laser power) for 90 bleaching iterations.

### Lentivirus production

Production of lentiviral particles was conducted using the second-generation packaging system as previously described [69, 70]. In brief, HEK293T cells were co-transfected with lentiviral plasmid shRNA and the packaging vectors pLP1, pLP2, and pLP-VSV-G (Thermo Fisher) using the Ca^2+^ phosphate transfection method. At 24, 36, and 48 h after transfection, the virus-containing supernatant was collected, and the lentiviral particles concentrated (800-fold) by ultracentrifugation (100,000 × g at 4 °C for 2 h) prior to aliquoting and storage at −80 °C.

### MT stability

WT and Mfn2 KO MEFs were incubated at 8 °C for 30 min to induce mild microtubule depolymerization. At the end of the incubation time, cells were gently washed with PEM 1× buffer (85 mM Pipes, pH 6.94, 10 mM EGTA, and 1 mM MgCl_2_) twice before extraction with PEM buffer, supplemented with 0.05 % Triton X-100. After 1 min extraction at 8 °C, a matching volume of fixative buffer (ice cold MetOH) was added dropwise to the coverslips, and cells were incubated for another 5 min at −20 °C. Cells were finally washed with PBS 1× and processed for immunofluorescence labeling. All images were analyzed using ImageJ software [71].

### Analysis of phospholipid synthesis in cultured cells

Both mitochondria and ER play key roles in the synthesis of phosphatidylserine (PtdSer), phosphatidylethanolamine (PtdEtn), and phosphatidylcholine (PtdCho). PtdSer is synthesized in the MAM; it then translocates to mitochondria, where it is converted to PtdEtn; PtdEtn then translocates back to the MAM, to generate PtdCho [72]. To test the effect of MFN2 on phospholipid synthesis mediated by MAM, WT and Mfn2 KO MEF cells were incubated for 2 h with serum-free medium to ensure removal of exogenous lipids. The medium was then replaced with MEM containing 2.5 μCi/ml of ^3^H-serine for 2, 4 and 6 h. The cells were washed and collected in DPBS, pelleted at 2500 g for 5 min at 4°C, and resuspended in 0.5 ml water, removing a small aliquot for protein quantification. Lipid extraction was done by the Bligh and Dyer method. Briefly, three volumes of chloroform/methanol 2:1 were added to the samples and vortexed. After centrifugation at 8000 g for 5 min, the organic phase was washed twice with two volumes of methanol/water 1:1, and the organic phase was blown to dryness under nitrogen. Dried lipids were resuspended in 60 μl of chloroform/methanol 2:1 (v/v) and applied to a TLC plate. Phospholipids were separated using two solvents, composed of petroleum ether/diethyl ether/acetic acid 84:15:1 (v/v/v) and chloroform/methanol/acetic acid/water 60:50:1:4 (v/v/v/v). Development was performed by exposure of the plate to iodine vapor. The spots corresponding to the relevant phospholipids (identified using co-migrating standards) were scraped and counted in a scintillation counter (Packard Tri-Carb 2900TR). Both mitochondria and ER play key roles in the synthesis of phosphatidylserine (PtdSer), phosphatidylethanolamine (PtdEtn), and phosphatidylcholine (PtdCho). PtdSer is synthesized in the MAM; it then translocates to mitochondria, where it is converted to PtdEtn; PtdEtn then translocates back to the MAM, to generate PtdCho [73]. Therefore, to test directly the effect of MFN2 mutations on phospholipid synthesis mediated by MAM, we incubated control and Mfn2 KO fibroblasts in medium containing ^3^H-serine and measured the incorporation of the label into newly-synthesized ^3^H-PtdSer and ^3^H-PtdEtn after 2 and 4 h.

### Lipidomics

All samples were collected and treated following recently accepted guidelines for the analysis of human blood plasma and/or serum. Lipids were extracted from equal amounts of material–(0.2 ml/sample) by a chloroform–methanol extraction method. Three comprehensive panels, scanning for either positive lipids, negative lipids or neutral lipids (under positive mode), were analyzed. Equal amounts of internal standards with known concentrations were spiked into each extract. Each standard was later used to calculate the concentrations of corresponding lipid classes by first calculating ratio between measured intensities of a lipid species and that of corresponding internal standard multiplied by the known concentration of the internal standard. Samples were analyzed using a 6490 Triple Quadrupole LC/MS system (Agilent Technologies, Santa Clara, CA). Cholesterol and cholesterol esters were separated with normal-phase HPLC using an Agilent Zorbax Rx-Sil column (inner diameter 2.1 Å~ 100 mm) under the following conditions: mobile phase A (chloroform:methanol:1 M ammonium hydroxide, 89.9:10:0.1, v/v/v) and mobile phase B (chloroform:methanol:water: ammonium hydroxide, 55:39.9:5:0.1, v/v/v/v); 95% A for 2 min, linear gradient to 30% A over 18 min and held for 3 min, and linear gradient to 95% A over 2 min and held for 6 min.

| INTERNAL STANDARD | Corresponding Lipid Class | Concentration (ug/ul) |
| --- | --- | --- |
| IS AcylPG 14:0-28:0 | Acyl PG, NAPE, NAPS | 0.046799614 |
| IS BMP 28:0 | BMP | 0.015298133 |
| IS CE C17 | CE | 78.59098931 |
| IS Cer C17:0 | Cer, dhCer | 0.758320608 |
| IS Chol d7 b | Free Cholesterol | 63.78791732 |
| **IS DG 4ME** | diacylglycerols | **0.640874053** |
| **IS dhSM**<br>**d18:0/12:0** | dihydrosphingomyelins | **2.579623778** |
| **IS DMPC** | AC | **12.34642208** |
| **IS GalCer**<br>**d18:1/12:0** | MhCer | **1.039897431** |
| **IS LacCer**<br>**d18:1/12:0** | LacCer | **0.259594347** |
| **IS LPC 13:0** | LPC | **12.34642208** |
| **IS LPE 14:0** | LPE | **0.098349468** |
| **IS LPI 13:0** | LPI | **0.07642123** |
| **IS MG C17** | MG | **0.242952978** |
| **IS PA 28:0** | PA | **0.068072997** |
| **IS PC 28:0** | PC | **12.34642208** |
| **IS PE 25:0** | PE | **8.839285714** |
| **IS PG 12:0/13:0** | PG | **0.446428571** |
| **IS PI 12:0/13:0** | PI | **2.232142857** |
| **IS PS 28:0** | PS | **11.92531331** |
| **IS SM d18:1/12:0** | SM | **13.39285714** |
| **IS Sulf d18:1/12:0** | Sulf | **0.225924621** |
| **IS TG 50:0 d5** | TG | **0.498018035** |

### Cellular fractionation

WT and Mfn2 KO fibroblasts were cultured on 15 cm petri dishes. Buffer A (10mM HEPES, 1.5 mM MgCl_2_, 10 mM KCl, 0.5 mM DTT, 0.05% NP40, pH 7.9) was prepared freshly and protease and phosphatase inhibitors were added. Cells were scraped thoroughly using buffer A and left on ice for 10 min. Samples were centrifuged at 3000 rpm for 10 min at 4°C and supernatants stored on ice. Pellets were resuspended in buffer B (5 mM HEPES, 1.5 mM MgCl_2_, 0.2 mM EDTA, 0.5 mM DTT, 26% Glycerol (v/v), pH 7.9) and added 4.6 M NaCl. Homogenize with 20 full stroke of Dounce homogenizer on ice and leave it on ice for 30 min.

### Isolation of adult DRG neurons

DRG were dissected from 8- to 10-wk-old C57BL/6J mice in cold Hank’s balanced salt solution (HBSS) (Life Technologies) or Dulbecco’s Modified Eagle’s medium (Life Technologies) and dissociated in 1 mg/mL Collagenase A for 1 h at 37 °C, followed by 0.05% trypsin (Life Technologies) digestion for 3 to 5 min at 37 °C and washed with Neurobasal medium (Invitrogen) supplemented with 2% B-27 (Invitrogen), 0.5 mM glutamine (Invitrogen), fetal bovine serum (FBS), and 100 U/mL penicillin-streptomycin. DRG neurons were then triturated by repeated gentle pipetting until no clump was visible, and neuronal bodies were resuspended in Neurobasal medium with FBS prior to plating onto 12 well plates (over 18 mm coverslips) that had been coated overnight with 100 μg/ mL poly-D-lysine at 37 °C and for 1 h at 37 °C with 10 μg/mL laminin (Life Technologies). After 30 min, Neurobasal medium, without FBS, was added to the plate. At 4 DIV, at least 30% of media was changed and 10 μM AraC was added to media every 4 d.

### Degeneration index in DRG neurons

As reported previously [69], images of an average of 10 random fields of dissociated adult DRG neurons fixed and immunostained with mouse anti-neurofilament antibody were acquired using a 20× objective lens (Olympus IX81) coupled to a monochrome CCD camera (Sensicam QE; Cooke Corporation). To quantify axonal degeneration, the areas occupied by the axons (total axonal area) and degenerating axons (fragmented axonal area) were measured in the same field from images of DRG neurons. Images were automatically thresholded (global threshold) using a default auto threshold method, binarized, and the fragmented axonal area measured by using the particle analyzer module of ImageJ (size of small fragments = 20 to 10,000 pixels). Degeneration index was calculated as the ratio between the fragmented axonal area and the total axonal area.

### Immunolabeling of *Drosophila* larvae

Immunolabeling of *Drosophila* larvae was performed largely as described previously [74]. Briefly, late third instar larvae were dissected in 1 × PBS, fixed in 4% paraformaldehyde (PFA, Electron Microscopy Sciences) in 1 × PBS for 15 min, washed three times in 1 × PBS + 0.3% Triton X-100 (PBS-TX), and blocked for 1h at room temperature (RT) or overnight at 4 °C in 5% normal donkey serum (NDS) in PBS-TX (Jackson Immunoresearch). Primary antibodies were chicken anti-GFP (1:1000; Abcam) and acetylated alpha-tubulin (1:400; Sigma Aldrich) diluted in 5% NDS in PBS-TX. The tissue was incubated overnight in primary antibodies at 4 °C and then washed in PBS-TX for 3 × 15 min at RT. Species-specific, fluorophore-conjugated secondary antibodies (Jackson ImmunoResearch) were used at 1:1000 in 5% NDS in PBS-TX and incubated overnight 4 °C. Tissue was washed in PBS-TX for 3 × 15 min. Immunolabeled tissue was mounted on poly-L-lysine coated coverslips, dehydrated 5 minutes each in an ascending ethanol series (30, 50, 70, 95, 2 × 100%), cleared in xylenes (2 × 10 min), and mounted in DPX (Fluka).

### Imaging and quantification of *Drosophila* sensory neurons

Images of somatosensory neurons from *Drosophila* larvae were acquired using a Yokogawa CSU-W1 SoRa mounted on a Zeiss Axio Observer using a 60× 1.46 NA Alpha Plan-Apochromat oil objective and a 4x magnification changer. Acquisitions included the cell body, axon, and dendrites of somatosensory neurons. Subsequent image analysis was performed using Fiji. Using the md neurons (109(80)2-Gal4, UAS-CD8-GFP) as reference, sub stacks covering the z-depth of each cell body were cropped and blinded for subsequent analysis. Additionally, 2-3 areas (300 × 300 px) devoid of neurons in the same image were selected to measure background levels of acetylated tubulin in each image and used to normalize the levels in the cell body. To measure acetylated tubulin levels in the cell bodies, cell bodies were selected using the polygon selection tool, and the area outside the cell body was cleared to avoid including acetylated tubulin staining surrounding the cell body in the subsequent quantification. Processed z-stacks of cell bodies were z-projected using average intensity. The mean gray value was measured and normalized against background levels of acetylated tubulin quantified in the same image. Raw images were used for quantification. Represented images shown in Fig. 5C were deconvolved using Microvolution (20 iterations).

### Co-immunoprecipitation assay

HEK293T cells were cultured in Dulbecco’s modified Eagle’s medium plus 10% fetal bovine serum (FBS), penicillin–streptomycin (1%) and L-glutamine (1%). Transient transfections were performed using DreamFect™Gold transfection reagent (Oz Biosciences SAS, Marseille, FR) in accordance with the manufacturer’s protocols. HEK293T cells were lysed in RIPA buffer (50 mM Tris-HCl at pH 7.6, 150 mM NaCl, 0.5% sodium deoxycholic, 5 mM EDTA, 0.1% SDS, 100 mM NaF, 2 mM NaPPi, 1% NP-40) supplemented with protease and phosphatase inhibitors, then centrifuged at 13,000 rpm for 30 min at 4 °C and the resulting supernatants were subjected to Bradford protein assay (Bio-Rad, Hercules, California, USA) for measuring total protein concentration. Immunoprecipitations were performed on 1mg of whole cell extracts by using anti-c-Myc agarose coniugated or anti-Flag M2 agarose (1–2 μg) for 2 h at 4 °C with rotation. c-Myc peptide or Flag-peptide (0,1 mg/ml) was used as a control. The immunoprecipitates were then washed five times with RIPA buffer, resuspended in sample loading buffer, boiled for 5 min, resolved in sodium dodecyl sulphate–polyacrylamide gel electrophoresis (SDS-PAGE), and then subjected to immunoblot analysis.

### Quantification and statistical analysis

Data are shown as median with interquartile range from at least 3 independent experiments and figures are generated by GraphPad Prism. Image analysis was performed by ImageJ (Fiji). Statistical analysis between two groups was performed Mann-Whitney *U* test and among 3 or more groups was performed using the Kruskal-Wallis with Dunn’s multiple comparisons test. Data are shown as means ±SEM, and statistical significance was analyzed by student’s t test (Table1) and two-way ANOVA with Dunnett’s multiple comparison (Table S1 and S2).

## Abbreviations

MT: Microtubule
PTMs: post-translational modifications
CMT2A: Charcot-Marie-Tooth Type 2A
MFN2: Mitofusin-2
ATAT1: α-tubulin acetyl transferase 1
HDAC6: Histone deacetylase 6
OMM: outer mitochondrial membrane
ER: endoplasmic reticulum
MFN1: Mitofusin-1
MAMs: mitochondria-associated ER membranes
TSA: trichostatin A
TTL: tubulin tyrosine ligase

## Acknowledgements

This study was funded by a TIGER Grant from the Taub Institute for Research on Alzheimer’s Disease and the Aging Brain at Columbia University to F.B., the RF1AG050658 (NIH/NIA) and R21NS120076 (NIH/NINDS) awards to F.B. We are grateful to Gregg G. Gundersen for stimulating discussions and access to his microscopes, and to Laurent Nguyen for sharing his ATAT1 constructs. Figures were prepared using PowerPoint and BioRender.

