## Supplementary Information for "MFN2-dependent recruitment of ATAT1 coordinates mitochondria motility with α-tubulin acetylation and is disrupted in CMT2A"

### MFN2-dependent recruitment of the ATAT1 coordinates mitochondria motility with α-tubulin acetylation and is disrupted in Charcot-Marie Tooth 2A

A. Kumar[1], D. Larrea[2], M.E. Pero[1,3], P. Infante[4], M. Conenna[4], G.J. Shin[5], W. B. Grueber[5,6], L. Di Marcotullio[4,7], E. Area-Gomez[2,8], and F. Bartolini[1,*]

[1]Department of Pathology & Cell Biology, Columbia University Irving Medical Center, 10032, New York, NY, USA

[2]Department of Neurology, Columbia University Irving Medical Center, 10032, New York, NY, USA

[3]Department of Veterinary Medicine and Animal Production, University of Naples Federico II, 80137, Naples, Italy

[4]Department of Molecular Medicine, 1st University of Rome "La Sapienza", 00161, Rome, Italy

[5]Department of Neuroscience, Zuckerman Mind Brain Behavior Institute, Columbia University, 10027, New York, NY, USA

[6]Department of Physiology & Cellular Biophysics, Zuckerman Mind Brain Behavior Institute, Columbia University, 10032, New York, NY, USA

[7]Istituto Pasteur-Fondazione Cenci Bolognetti, University of Rome La Sapienza, Rome, Italy.

[8]Current address: Department de Biología Celular y Molecular, Centro de Investigaciones Biológicas, MARGARITA SALAS, CSIC, 28040, Madrid, Spain

#### List of supplementary information:

**Supplementary Figures**
Figure S1
Figure S2
Figure S3
Figure S4
Figure S5

**Supplementary Tables**
Table S1
Table S2

Figure S1

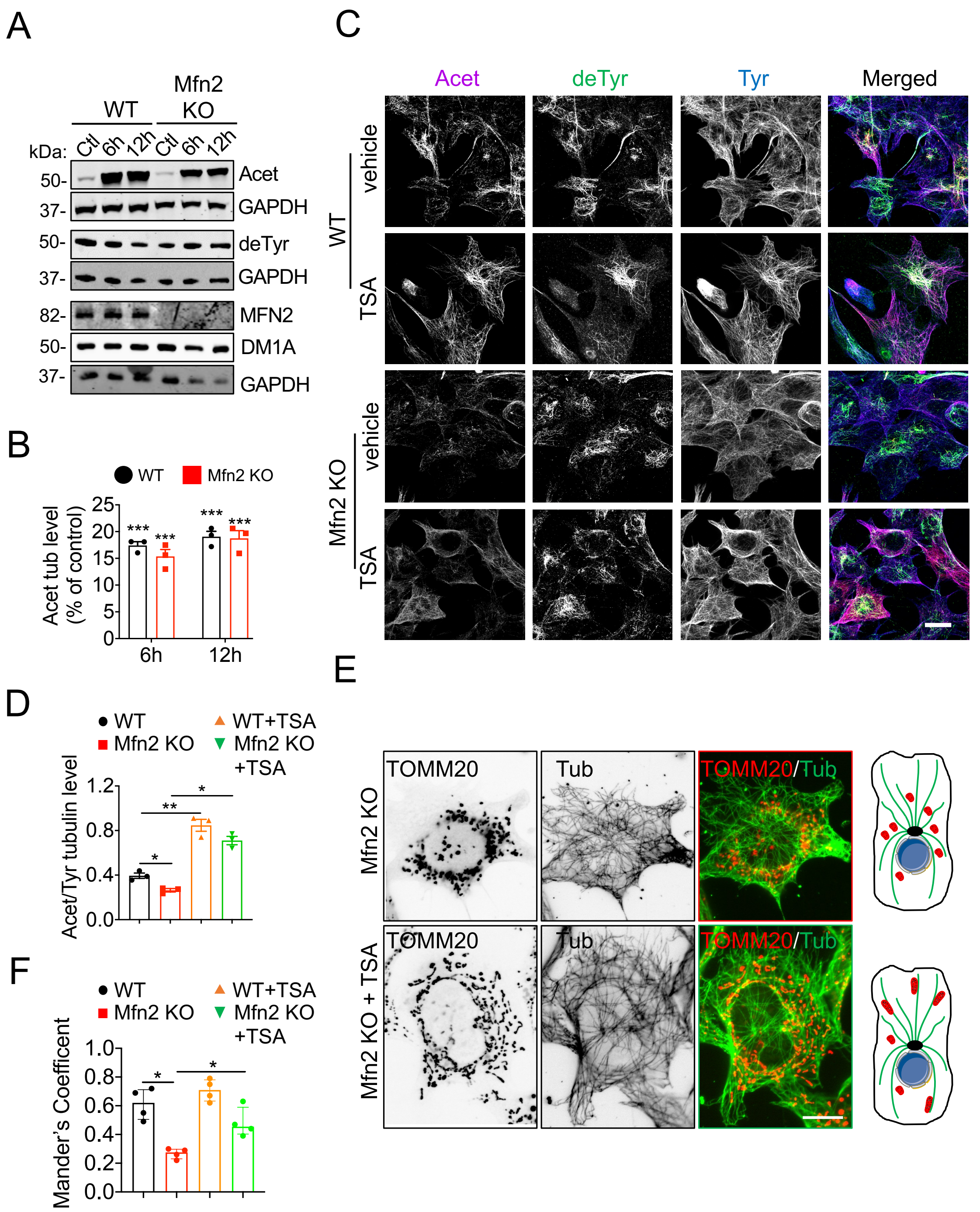


**Figure S1. HDAC6 inhibition restores acetylated tubulin levels and mitochondria association to MTs in Mfn2 KO MEFs.** (A) Representative immunoblot of acetylated (Acet), detyrosinated (deTyr), total α-tubulin (DM1A), mitofusin-2 (MFN2) and GAPDH levels in whole cell lysates from WT and Mfn2 KO MEFs incubated with control vehicle or 10 µM of trichostatin A (TSA) for 6 h and 12 h prior to lysis. (B) Quantification of acetylated tubulin relative to control levels (%) in WT and Mfn2 KO MEFs treated as in A. (C) Representative immunofluorescence staining of acetylated (Acet), detyrosinated (deTyr) and tyrosinated (Tyr) tubulin in WT and Mfn2 KO MEFs incubated with control vehicle or TSA for 6 h. Scale bar, 10 µm. (D) Quantification of acetylated to tyrosinated tubulin (bulk tubulin marker) immunofluorescence signal ratio measured in individual cells treated as in C. Scale bar, 10 µm. (E) Representative immunofluorescence staining of mitochondria and tyrosinated tubulin (Tub) in Mfn2 KO MEFs incubated with control vehicle or 10 nM of TSA for 6 h. Scale bar, 5 µm. (F) Quantification of mitochondria associated with MTs (identified by the bulk tubulin marker tyrosinated tubulin) using Mander's coefficient. Data are expressed as median with interquartile range from 3 independent experiments. * $p<0.05$; ** $p<0.01$; *** $p<0.001$; ns non-significant by Kruskal-Wallis test.

Figure S2

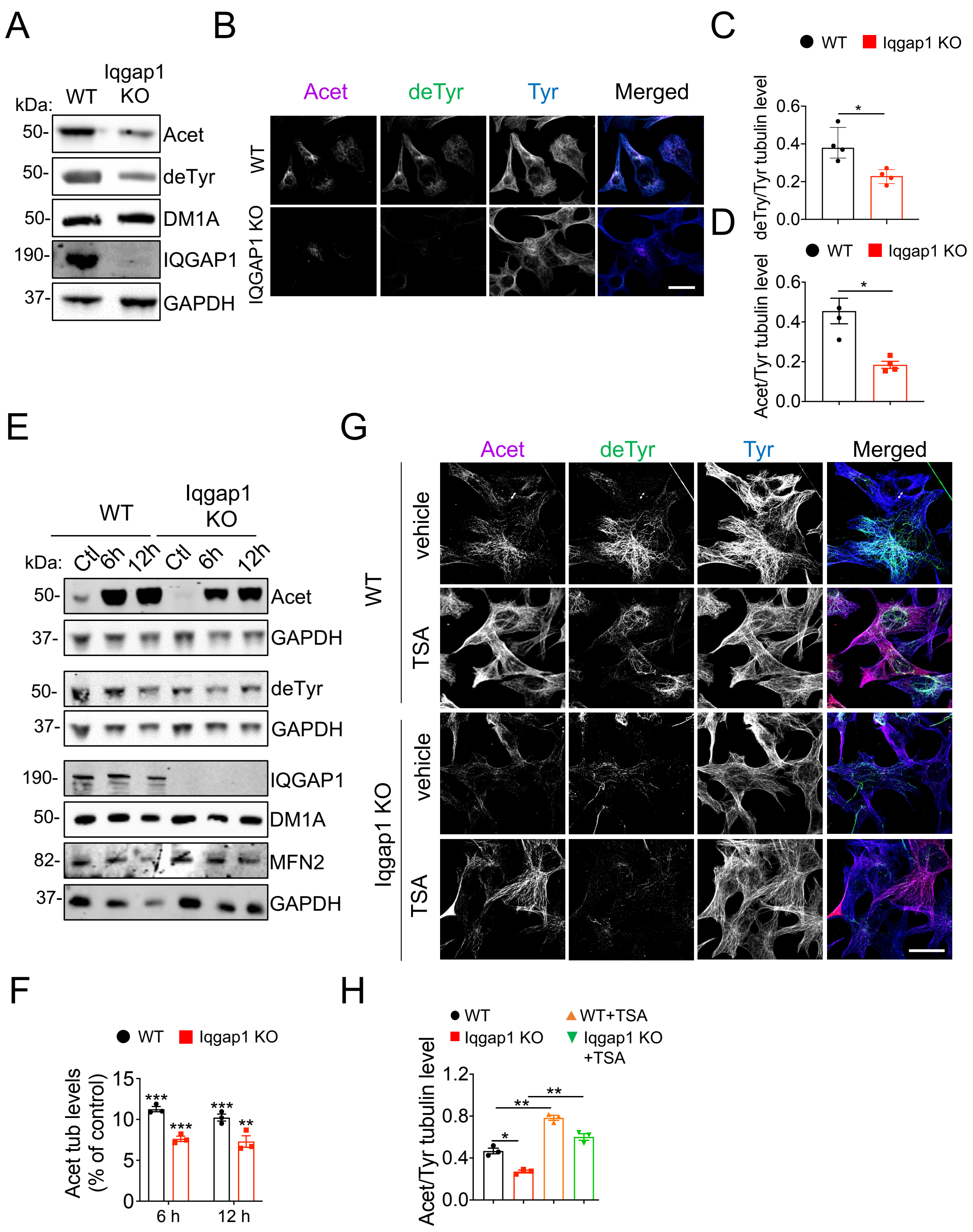


**Figure S2**. **HDAC6 inhibition restores acetylated α-tubulin levels in Iqgap1 KO MEFs**. (A) Representative immunoblot of WT and Iqgap1 KO whole MEF lysates. DM1A, total α-tubulin, Acet, acetylated tubulin, deTyr, detyrosinated tubulin. GAPDH, loading control. (B) Representative immunofluorescence staining (images are maximum projections from z-stacks) of acetylated (Acet), detyrosinated (deTyr) and tyrosinated (Tyr) tubulins in WT and Iqgap1 KO MEFs. (C) Quantification of detyrosinated tubulin signal in Iqgap1 KO MEFs relative to WT levels. (D) Quantification of acetylated tubulin signal in Iqgap1 KO MEFs relative to WT levels. (E) Representative immunoblot of acetylated (Acet), detyrosinated (deTyr), total (DM1A) α-tubulin, IQGAP1, mitofusin-2 (MFN2) and GAPDH levels in WT and Iqgap1 KO MEFs incubated with 10 nM of the HDAC6 inhibitor trichostatin A (TSA) or vehicle control for 6 h or 12 h prior to lysis. (F) Quantification of acetylated tubulin (Acet) levels relative to vehicle control (%) in WT and Iqgap1 KO MEFs incubated with 10 nM TSA for 6 or 12 h. (G) Representative immunofluorescence staining (images are maximum projections from z-stacks) of WT and Iqgap1 KO MEFs incubated with 10 nM of TSA or vehicle control for 6 h prior to fixation. (H) Quantification of acetylated (Acet) to tyrosinated (Tyr) tubulin immunofluorescence signal ratio measured in WT and Iqgap1 KO MEFs as in G. Data are median with interquartile range. *, $p \leq 0.05$; **, $p \leq 0.01$; ***, $p \leq 0.001$; ****, $p \leq 0.0001$; ns non-significant by Mann–Whitney U test (S3C and D) or Kruskal-Wallis test (S3F and H). (n=3-4 independent experiments); Scale bar, 10 μm.

Figure S3

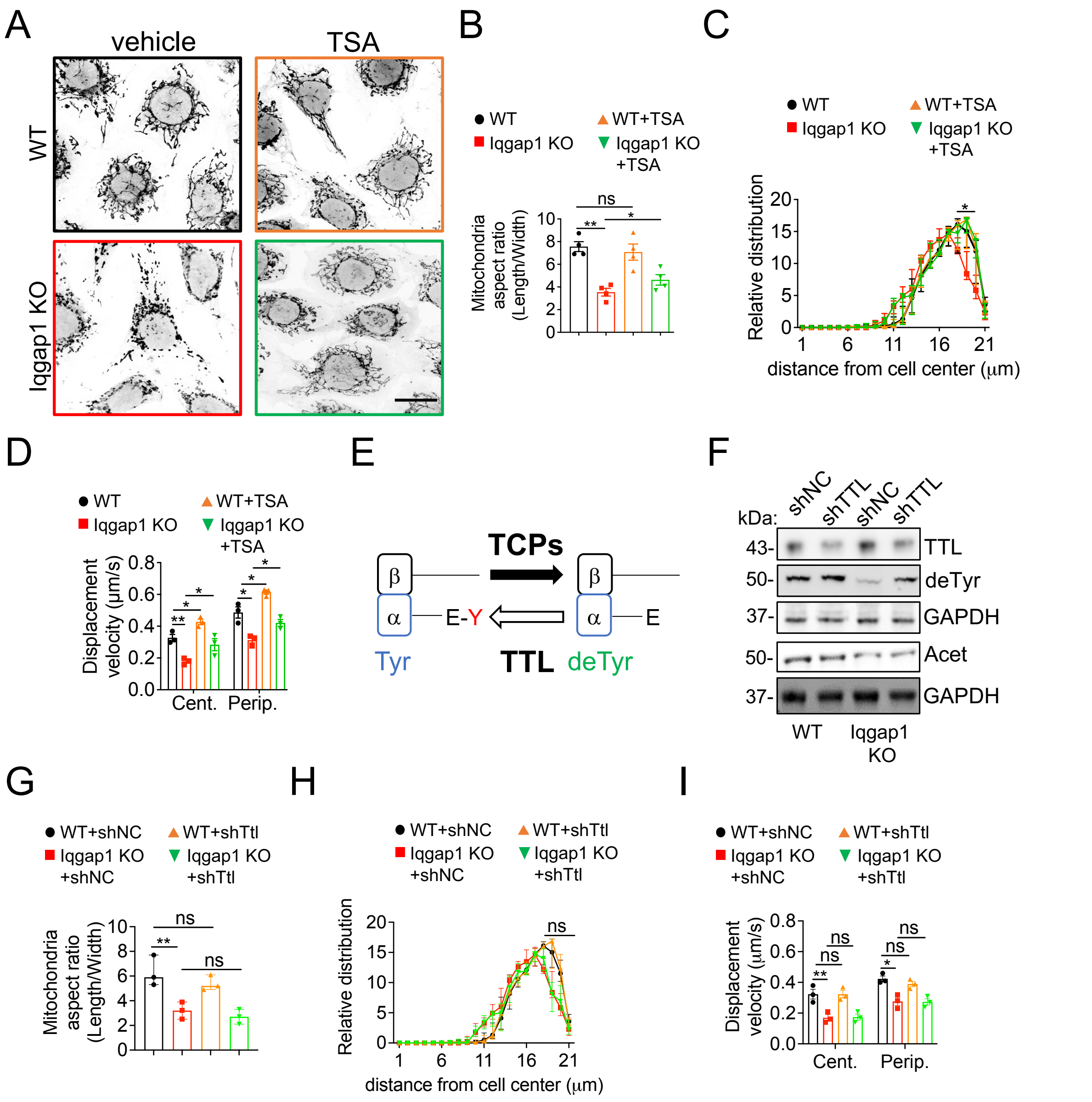


**Figure S3**. **Restoring α-tubulin acetylation levels rescues mitochondria dynamics in Iqgap1 KO MEFs.** (A) WT and Iqgap1 KO control and trichostatin A (TSA) treated (10 nM, 6 h) MEFs were stained with mitoTracker Red. (B) Mitochondria morphology is calculated as ratio of length /width in cells treated as in A. (C) Relative distribution of mitochondria from the cell center in MEFs treated as in A. (D) Mitochondrial displacement velocity near the cell center and periphery in MEFs as in A. Live-imaging was performed for 3 min (1f/2s). (E) Schematic of the enzymes involved in the detyrosination/retyrosination α-tubulin cycle. Note that detyrosination of α-tubulin occurs on MTs while α-tubulin retyrosination occurs on the soluble tubulin heterodimer. TCPs, tubulin carboxypeptidases; deTyr, detyrosinated α-tubulin; Tyr, tyrosinated α-tubulin. TTL, tubulin tyrosine ligase. (F) Representative immunoblot of whole cell lysates from WT and Iqgap1 KO MEFs depleted of tubulin tyrosine ligase (TTL) expression by shRNA transfection. TTL, tubulin tyrosine ligase; DeTyr, detyrosinated tubulin; Acet, acetylated tubulin; GAPDH, loading control. (G) Mitochondria morphology is calculated as ratio of length/width in cells depleted of TTL expression. (H) Relative distribution of mitochondria from the cell center in MEFs depleted of TTL expression. (I) Mitochondrial displacement velocity near the cell center and periphery in MEFs depleted of TTL expression. Live-imaging was performed for 3 min (1f/2s). Data are expressed as median with interquartile range n = 150-250 mitochondria from 15-30 cells in 3-4 independent exp. *, $p \leq 0.05$; **, $p \leq 0.01$; ns non-significant by Kruskal-Wallis test. Scale bar, 10 μm.

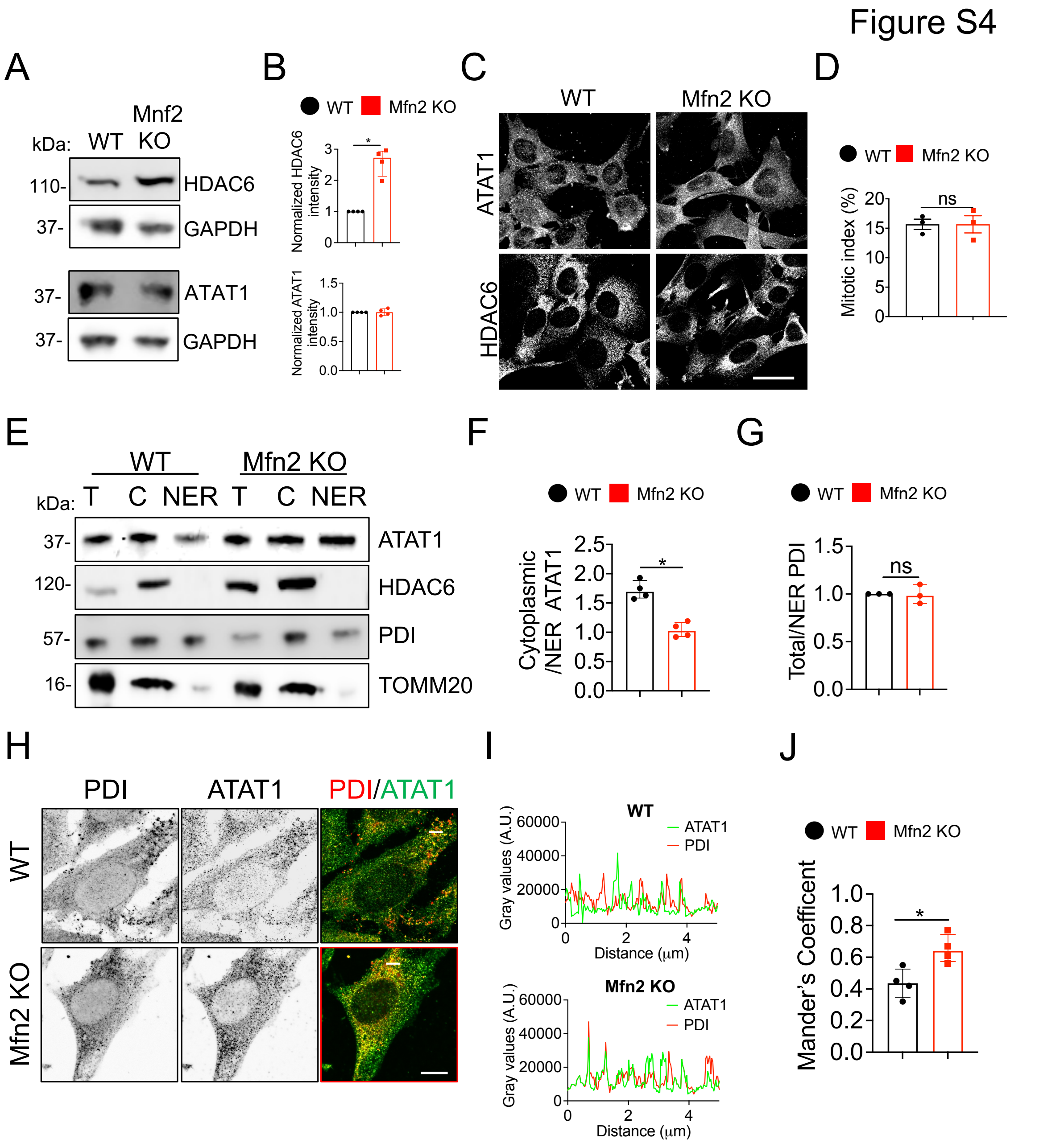


**Figure S4. MFN2 regulates HDAC6 levels and localization of ATAT1 to intracellular membranes.** (A) Representative immunoblot of HDAC6 and ATAT1 levels in WT and Mfn2 KO whole MEF lysates. GAPDH, loading control. (B) Quantification of expression level of HDAC6 and ATAT1 in Mfn2 KO MEFs relative to WT MEFs. (C) Representative immunofluorescence staining of ATAT1 and HDAC6 in WT and Mfn2 KO MEFs. (n=200-225 cells). Scale bar, 10 µm. (D) Mitotic index assessed by nuclear DAPI staining in WT and Mfn2 KO MEFs. (n=150 cells). (E) Representative immunoblot of cytosolic and nuclear envelope/endoplasmic reticulum (ER) fractions (T=total lysate; C=cytoplasmic fraction; NER= nuclear ER fraction) from WT and Mfn2

KO MEF lysates. (F) Quantification of ATAT1 levels in cytoplasmic versus NER fraction expressed as ratio of intensity values from analysis as in E. (G) Quantification of the ER marker PDI levels in total versus NER fraction expressed as ratio of intensity values. (H) Max projection confocal image of WT and Mfn2 KO cells stained with PDI and ATAT1 (n=20-25 cells). Scale bar, 10 μm. (I) Line scan analysis of PDI and ATAT1 from selected regions (white bars in H) in WT and Mfn2 KO cells. (J) Quantification of localization of ATAT1 and PDI in WT and Mfn2 KO MEFs by Mander's coefficient. Data are median with interquartile range from 3-4 independent experiments. * $p \leq 0.05$, ** $p < 0.01$; ns non-significant by Mann-Whitney U test.

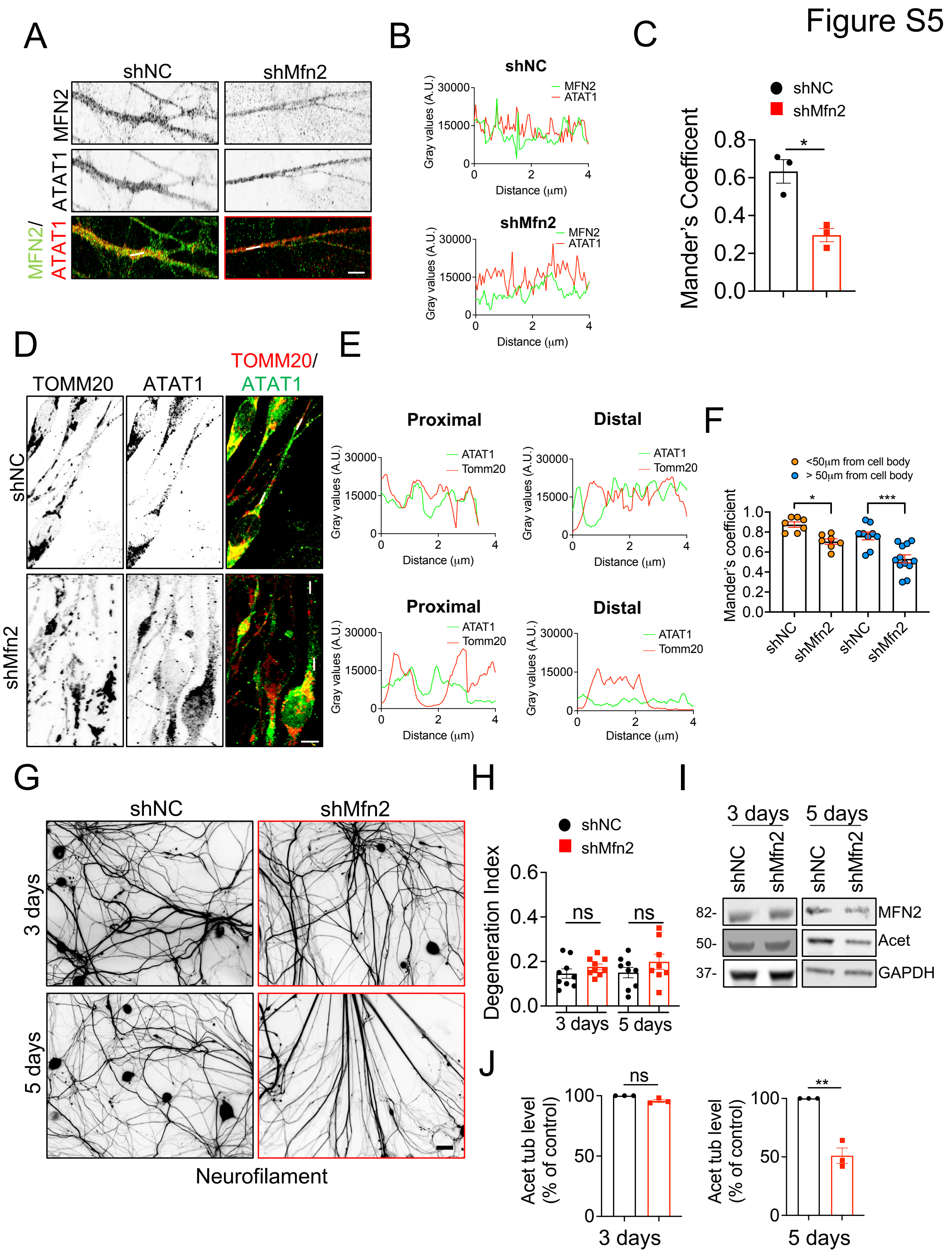


**Figure S5. Loss of acetylated tubulin precedes axonal degeneration in DRG neurons deprived of MNF2.** (A) Airyscan confocal images of ATAT1 and MFN2 in shNC and shMfn2 knockdown adult DRG neurons (14 days) silenced of Mfn2 expression at 7 DIV. (B) Linescan analysis of ATAT1 and MFN2 from selected regions (white bars in A) in shNC and shMfn2 knockdown adult DRG neurons (14 days) silenced of Mfn2 expression at 7 DIV. (C) Quantification of localization of MFN2 and ATAT1 in shNC and shMfn2 knockdown adult DRG neurons (14 days) silenced of Mfn2 expression at 7 DIV by Mander's correlation coefficient. (D) Confocal images of staining of TOMM20 (red) and ATAT1 (green) in shNC and shMfn2 knockdown adult DRG neurons (14 days) silenced of Mfn2 expression at 7 DIV. (E) Line scan analysis at proximal and distal neurites from selected regions (white bars in D) in shNC and shMfn2 knockdown adult DRG neurons (14 days) silenced of Mfn2 expression at 7 DIV. (F) Mander's coefficient analysis of TOMM20 and ATAT1 in shNC and shMfn2 knockdown adult DRG neurons (14 days) silenced of Mfn2 expression at 7 DIV. (G) Neurofilament staining of shNC and shMfn2 knockdown DRG neurons silenced of Mfn2 expression for 3 days and 5 days at 7 DIV. Scale bar, 50 µm. (H) Degeneration index of axons in shNC and shMfn2 kd DRG neurons silenced for Mfn2 expression for 3 days and 5 days at 7 DIV. (I) Representative immunoblot of MFN2 and acetylated tubulin (Acet) levels in DRG neurons silenced of Mfn2 expression for 3 days and 5 days at 7 DIV. GAPDH, loading control. Data are expressed as median with interquartile range. n= 20-25 cells from 3 independent experiments. * $p \leq 0.05$, ** $p<0.01$; *** $p<0.001$ ns: non-significant by Mann–Whitney U test. Scale bars, 10 µm.

#### Table S1

| | WT | Mfn2 KO | WT+ TSA | Mfn2 KO + TSA |
|---|---|---|---|---|
| Growth rate (µm/s) | 0.06 ± 0.01 | 0.11 ± 0.008 * | 0.04 ± 0.007 | 0.08 ± 0.005 * |
| Shrinkage rate (µm/s) | 0.08 ± 0.004 | 0.11 ± 0.009 * | 0.07 ± 0.008 | 0.08 ± 0.005 * |
| Catastrophe freq. (s-1) | 0.06 ± 0.006 | 0.06 ± 0.006 | 0.05 ± 0.004 | 0.05 ± 0.006 |
| Rescue freq. (s-1) | 0.08 ± 0.006 | 0.08 ± 0.006 | 0.07 ± 0.006 | 0.09 ± 0.004 |
| % Growth | 39.5 ± 1.32 | 49.95 ± 0.95 *** | 20.04 ± 1.28 **** | 22.4 ± 0.90 **** |
| % Shrinkage | 32.05 ± 0.87 | 37.68 ± 1.77 * | 18.25 ± 1.37 **** | 16.05 ± 1.42 **** |
| % Pause | 20.68 ± 0.71 | 17.1 ± 1.37 | 63.5 ± 0.64 **** | 62.13 ± 1.68 **** |
| MT lifetime (s) | 60.25 ± 2.05 | 61.5 ± 1.93 | 56.5 ± 3.66 | 60.25 ± 1.43 |
| MT dynamicity (µm/min) | 6.6 ± 0.64 | 11.63 ± 0.43 *** | 4.17 ± 0.44 * | 4.858 ± 0.46 **** |
| Number of MTs | 22 | 22 | 24 | 24 |

**Table S1. HDAC6 inhibition normalizes MT dynamics in Mfn2 KO MEFs.** MT dynamics were measured from time-lapse analysis of GFP-tubulin-labeled MTs using epifluorescence microscopy in WT and Mfn2 KO MEFs treated with 10 nM trichostatin A (TSA) for 6 h. Live imaging of MT dynamics was performed for 5 min (1f/5s). * in black represents statistical comparison with WT control while * in red represents statistical comparison with Mfn2 KO control. Data are mean ± SEM from 3 independent experiments. * $p<0.05$; ** $p<0.01$; *** $p<0.001$; **** $p<<0.001$ by 2-way ANOVA with Dunnett's multiple comparison.

Table S2

| | WT | Iqgap1 KO | WT+ TSA | Iqgap1 KO + TSA |
|---|---|---|---|---|
| Growth rate (µm/s) | 0.06 ± 0.008 | 0.13 ± 0.004 *** | 0.04 ± 0.004 | 0.095 ± 0.006 ** |
| Shrinkage rate (µm/s) | 0.04 ± 0.010 | 0.12 ± 0.006 *** | 0.03 ± 0.004 | 0.09 ± 0.004 |
| Catastrophe freq. (s-1) | 0.07 ± 0.004 | 0.06 ± 0.007 | 0.06 ± 0.003 | 0.06 ± 0.004 |
| Rescue freq. (s-1) | 0.07 ± 0.006 | 0.08 ± 0.010 | 0.07 ± 0.007 | 0.08 ± 0.006 |
| % Growth | 45.5 ± 2.021 | 45 ± 1.354 | 23.75 ± 0.85 *** | 25.25 ± 1.54 *** |
| % Shrinkage | 31.5 ± 2.63 | 30.18 ± 2.254 | 27.25 ± 3.19 | 22.5 ± 0.86 *** |
| % Pause | 22.5 ± 1.041 | 24.75 ± 1.652 | 50 ± 2.48 *** | 51.75 ± 1.10 *** |
| MT lifetime (s) | 47.5 ± 2.533 | 51 ± 1.472 | 49.5 ± 2.32 | 51.25 ± 1.79 |
| MT dynamicity (µm/min) | 5.22 ± 0.317 | 8.17 ± 0.606 ** | 3.32 ± 0.24 ** | 3.77 ± 0.33 ** |
| Number of MTs | 20 | 21 | 25 | 25 |

**Table S2**. **HDAC6 inhibition rescues MT dynamics in Iqgap1 KO MEFs**. MT dynamics were measured from time-lapse analysis of WT and Iqgap1 KO MEFs transfected with EGFP-tubulin prior to treatment with vehicle or 10 nM trichostatin A (TSA) for 6 h. Live imaging was performed for 5 min (1f/5s). * black represents statistical comparison with WT control and * red represents statistical comparison with Iqgap1 KO control. Data are mean ± SEM from 3 independent experiments. * $p<0.05$; ** $p<0.01$; *** $p<0.001$; by 2-way ANOVA with Dunnett's multiple comparison.
